# Generating functional and multistate proteins with a multimodal diffusion transformer

**DOI:** 10.1101/2025.09.03.672144

**Authors:** Bowen Jing, Anna Sappington, Mihir Bafna, Ravi Shah, Adrina Tang, Rohith Krishna, Adam Klivans, Daniel J. Diaz, Bonnie Berger

**Author notes:** Equal contribution (Bowen Jing, Anna Sappington). **Code:** https://github.com/bjing2016/ProDiT.

## Abstract

Generating proteins with the full diversity and complexity of functions found in nature is a grand challenge in protein design. Here, we present ProDiT, a multimodal diffusion model that unifies sequence and structure modeling paradigms to enable the design of functional proteins at scale. Trained on sequences, 3D structures, and annotations for 214M proteins across the evolutionary landscape, ProDiT generates diverse, novel proteins that preserve known active and binding site motifs and can be successfully conditioned on a wide range of molecular functions, spanning 465 Gene Ontology terms. We introduce a diffusion sampling protocol to design proteins with multiple functional states, and demonstrate this protocol by scaffolding enzymatic active sites from carbonic anhydrase and lysozyme to be allosterically deactivated by a calcium effector. Our results showcase ProDiT’s unique capacity to satisfy design specifications inaccessible to existing generative models, thereby expanding the protein design toolkit.

## 1 Introduction

Generative models trained on datasets of protein sequences or backbone structures have enabled significant advances in *de novo* protein design [11, 51]. Recently, structure diffusion models [49, 22, 26, 2] have been developed to generate protein binders [49, 26] or scaffold functional motifs [49, 2] while fine-tuned protein language models [30, 14, 4, 31, 18, 8] have been prompted to generate functional sequences with low homology to existing ones [30, 36, 18]. However, these existing paradigms focus only on structure or sequence in isolation and are therefore fundamentally incapable of capturing the complete functional landscape of natural proteins. Structure generative models require hand-crafted specification of function in terms of a binding partner or functional motif [49], depend on slow and specialized architectures adopted from protein structure prediction [6], and are limited to the relatively scarce structural training data in the Protein Data Bank (PDB). On the other hand, sequence generative models cannot natively reason about sequence-structure-function relationships, have shown lower generation quality compared to structure generative models [4], and have relied on fine-tuning [30] or complex prompting and filtering [18] to generate functional proteins. Neither family of approaches can target protein *dynamics*, a key feature of natural proteins, which often adopt multiple states to modulate their functions (e.g., motor proteins, allosterically regulated enzymes, signaling proteins). We hypothesize that a generative model, efficiently trained at scale with a broad understanding of protein functional space via both sequence and structure, would unlock many crucial protein design capabilities.

Here, we present ProDiT (Protein Diffusion Transformer), a novel framework that unifies structural and sequence generative modeling into a multimodal diffusion generative process, to enable these design capabilities. ProDiT leverages direct diffusion-based modeling of 3D structural coordinates alongside amino acid tokens using a fast, scalable transformer architecture, eschewing the specialized architectures typically used for modeling protein structure [6]. Our model design allows scaling multimodal training across large protein datasets, expanding beyond the PDB to the 214M available structures in AlphaFoldDB [44] and utilizing the full set of functional annotations in UniProtKB. Compared to previous attempts at multimodal protein generative modeling such as ProteinGenerator [29] and ESM3 [18], ProDiT models sequence and structure with diffusion processes in their respective state spaces, instead of tokenizing structure into a *discrete* vocabulary (ESM3) or forcing sequence generation into a *continuous* diffusion framework (ProteinGenerator). These prior approaches exhibit significantly degraded generation quality compared to models that specialize in generating structure, such as RFDiffusion [49]. In contrast, our model matches or often exceeds the unconditional structure generation quality of RFDiffusion across all sequence lengths, while also significantly exceeding the sequence generation quality of dedicated protein language models (PLMs) like EvoDiff [4].

We explore ProDiT’s broad understanding of the protein sequence and structural landscape by conditioning generation on functional descriptors in the form of molecular function Gene Ontology (GO) terms. Although prior works have included similar annotation inputs in their training [30, 18], they do not report detailed evaluations of function-conditioned generation. We conduct comprehensive benchmarking of ProDiT across 915 diverse GO terms representing highly specific descriptions of protein function and find generations predicted as functional for 463 of these terms, including terms represented in less than 0.01% of training examples in UniProtKB. Many of these generations have only moderate structural similarity to known functional proteins (TM-score *<* 0.7), showing an ability to generalize across the structural landscape. We also build upon techniques from image generative modeling for prompt adherence to further boost GO term adherence, a family of techniques not applicable to tokenized models such as ESM3. Structural alignments of our generations against known functional proteins reveal atomic-level recovery of active site residues, despite low global sequence similarity. Our designs additionally recover binding residues to enzymatic cofactors, e.g., NAD(P)+ for aldehyde dehydrogenase and CoA for malonyltransferase, demonstrating a multifaceted understanding of protein function.

Leveraging ProDiT’s multimodal capabilities, we also develop a principled protocol for the design of dynamic, multistate proteins, derived from a novel interpretation of the sequence-structure-function paradigm based on probabilistic graphical models. When distinct motifs are provided for the desired output protein states, our protocol generates two separate structural scaffolds coupled to a common sequence in a single generative rollout. As such, our approach represents a significant advance over prior protocols requiring brute-force sampling of plausible alternative conformations, previously thought necessary for such a protein design task [35, 17]. We demonstrate our approach by scaffolding active sites from carbonic anhydrase and lysozyme to be allosterically modulated by calcium binding, an important step towards the design of custom and controllable enzymes.

## 2 Results

### 2.1 Multimodal Generation of Sequence and Structure

We sought to develop a general framework that combines methods from protein language models and structure diffusion models to enable multimodal generation. To do so, ProDiT iteratively reverses two diffusion processes: one consisting of Gaussian noise added to the C*α* coordinates of centered protein structures and the other consisting of random masking of the amino acid sequence (Fig. 1A; **Methods**). At training time, the noise levels for sequence and structure are sampled independently; as a result, the model is able to denoise these modalities asynchronously, allowing for multiple generation workflows: sequence generation, structure generation (optionally followed by inverse folding), or sequence and structure co-generation. Additionally, ProDiT can accept as input structural or functional constraints in the form of motifs or molecular function Gene Ontology (GO) terms, respectively, for conditional generation. The model is trained and sampled with current best practices from the discrete and continuous diffusion literature (**Methods**).

**Figure 1:**
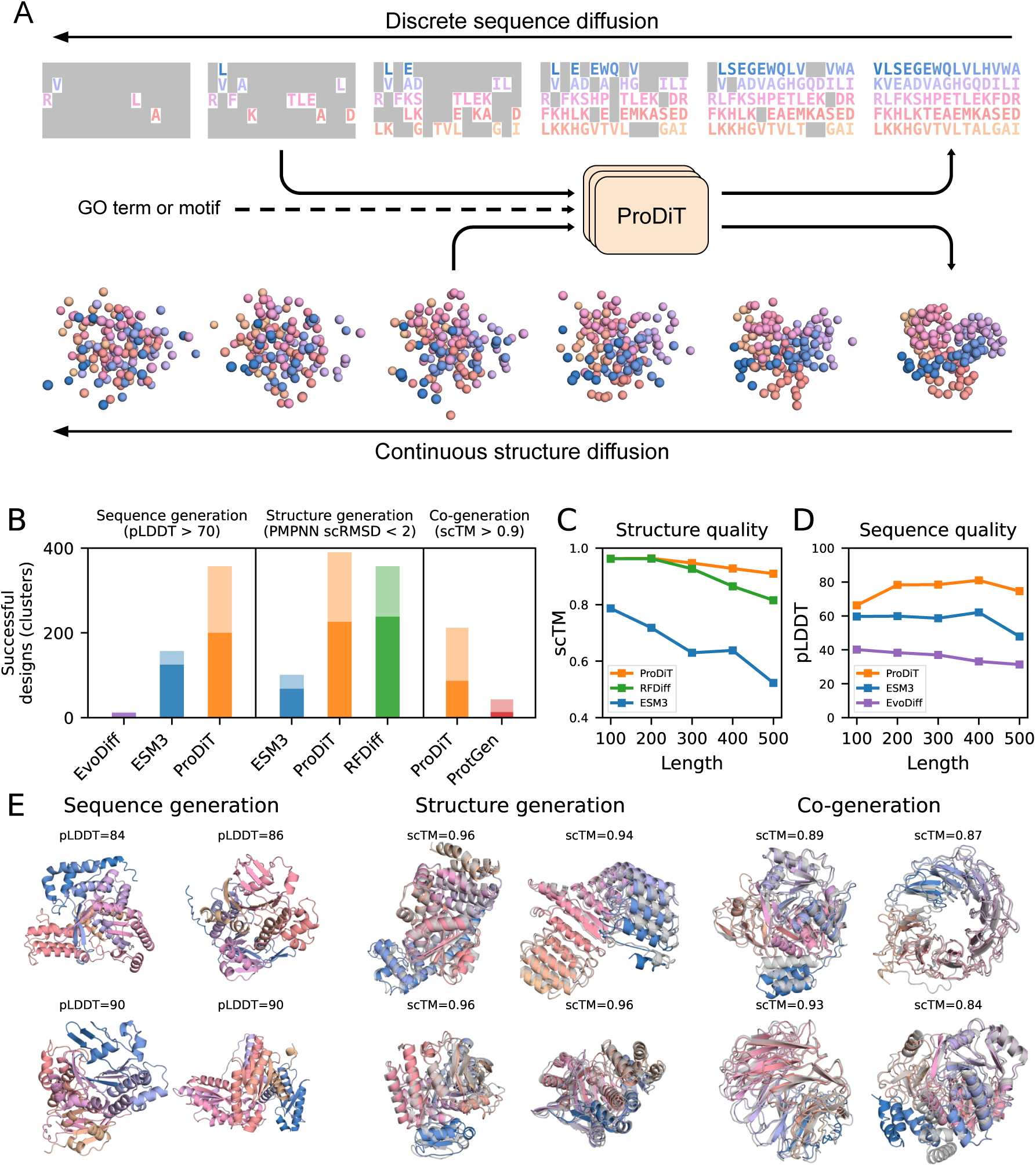
Overview of ProDiT and unconditional generation. **(A)** ProDiT is trained to denoise a joint diffusion process over sequence and structure. The two inputs may be at different noise levels, allowing for flexible generation workflows. Optionally, ProDiT can also condition on a GO term or structural motif. **(B)** Comparison of ProDiT with other models by number of successful generations (light color) and unique FoldSeek clusters (dark color) from 500 samples (100 each of length 100, 200, 300, 400, 500). **(C)** Structure generation quality across protein lengths, measured in terms of self-consistency TM-score (scTM) between the generated structure and ProteinMPNN-designed sequence. ProDiT maintains structure quality better than other methods at longer sequence lengths. **(D)** Sequence generation quality across protein lengths, measured in terms of ESMFold predicted Local Distance Difference Test (pLDDT). ProDiT obtains much higher sequence quality than other methods at all lengths. **(E)** Selected samples from ProDiT of length 500 under each generation protocol, labeled with the corresponding pLDDT or scTM metrics. For structure- and co-generation, the ESMFold refolded structure is superimposed in grey. RFDiff: RFDiffusion; ProtGen: ProteinGenerator.

We first assess the performance of ProDiT in unconditional generation of sequence and structure according to standard metrics (Fig. 1B–D). For each method and modality, we draw 100 samples each of lengths 100, 200, 300, 400, and 500 amino acids and compute the total number of successful generations. For generated sequences, success is defined by predicted Local Distance Difference Test (pLDDT) *>* 70 after folding with ESMFold [28]. For generated structures, we produce a corresponding sequence with ProteinMPNN and refold with ESMFold; success is defined by self-consistency RMSD (scRMSD) *<* 2Å. We assess diversity by clustering generated structures using FoldSeek [43] and report the number of unique clusters. For all generation modalities, we visualize proteins of length 500 amino acids sampled from ProDiT and observe diverse secondary structure elements, varied topologies, and globular, symmetric, and compact geometries (Fig. 1E).

ProDiT generates significantly more successful sequences than ESM3 and EvoDiff across all sequence lengths, surpasses the structure generation success rate of ESM3 [18], and matches the performance of RFDiffusion, a well-known structure generative model [49]. The difference in structure generation quality is most substantial at longer sequence lengths (*>* 300 amino acids) (Fig. 1C), where ESM3 protein generations become highly unstructured and RFDiffusion quality deteriorates more rapidly than ProDiT, potentially due to the larger training crops (512 vs 256) enabled by our simpler architecture. We observe similar trends when computing generation quality in terms of mean pLDDT or self-consistency TM-score (scTM) for sequence and structure, respectively (Fig. S1,S2). Strikingly, ProDiT accomplishes all this with a 5x smaller model than ESM3 (321M vs 1.7B parameters) and without using any equivariant modules or 2D tracks in its architecture.

For co-generation, ProDiT produces self-consistent sequence and structure pairs when simultaneously denoising over both modalities (Fig. 1B, right), using a success cutoff of scTM *>* 0.9 after refolding with ESMFold. A previously reported approach for co-generation, ProteinGenerator [29], adopts the architecture of RFDiffusion while embedding protein sequence into continuous space in order to apply continuous diffusion. However, ProteinGenerator is largely unable to generate self-consistent designs at lengths *>* 200 amino acids, whereas ProDiT’s multimodal diffusion can successfully generate self-consistent designs at 500 amino acids (Fig. S3). To further investigate ProDiT’s ability to generate self-consistent proteins, we used ProDiT to generate a backbone and then to inverse fold a corresponding sequence, demonstrating its ability to reason over both modalities successively (Fig. S5). This capability is crucial for function-conditioned generation, as functional properties are determined by both primary sequence and backbone structure. In contrast, inverse folding models that predict sequences solely based on backbone structures (like ProteinMPNN) lack an intrinsic understanding of whether those sequences will possess a desired function.

### 2.2 Steering Design with Molecular Function

We hypothesized that a model with broad understanding of sequence and structure across the evolutionary landscape could be steered towards generating diverse proteins with specific functions. To demonstrate this capability, we assessed the responsiveness of ProDiT to functional conditioning with molecular function Gene Ontology (GO) terms. To systematically screen functional designs *in silico*, we employ the DeepFRI function prediction network [16] to predict a probability for each GO term given an input structure. In total, 915 unique molecular function GO terms are present in both the input vocabulary of ProDiT and the output vocabulary of DeepFRI (Tab. S1); each of these terms is associated with a median of 0.08% of proteins in UniProtKB (0.03%–0.26% IQR), providing a highly specific description of protein function. For each of these GO terms, we generate 100 samples of length 300 amino acids from ProDiT (first generating the structure, then the sequence) and re-fold the sequence with ESMFold (**Methods**). A generation is counted as successful if the corresponding GO term is assigned by DeepFRI with *>* 50% confidence for the refolded structure.

#### 2.2.1 Systematic Assessment of Function Conditioning

In aggregate, 463 GO terms have at least one successful ProDiT design, with success rates being higher for the most prevalent GO terms and lower for less prevalent ones (Fig. 2A,B), suggesting ProDiT function-conditioned generation is data limited. We assess the diversity of the generations by computing the TM-scores between successful designs with the same GO term and novelty via the TM-score to the most similar structure in AlphaFoldDB with the same GO term (Fig. 2C). We observe slightly improved diversity for more common GO terms versus less common terms (0.60 vs 0.66 TM-score, respectively), suggesting more robust generalization for frequently observed terms, although the novelty is similar (0.81 vs 0.78 TM-score, respectively).

**Figure 2:**
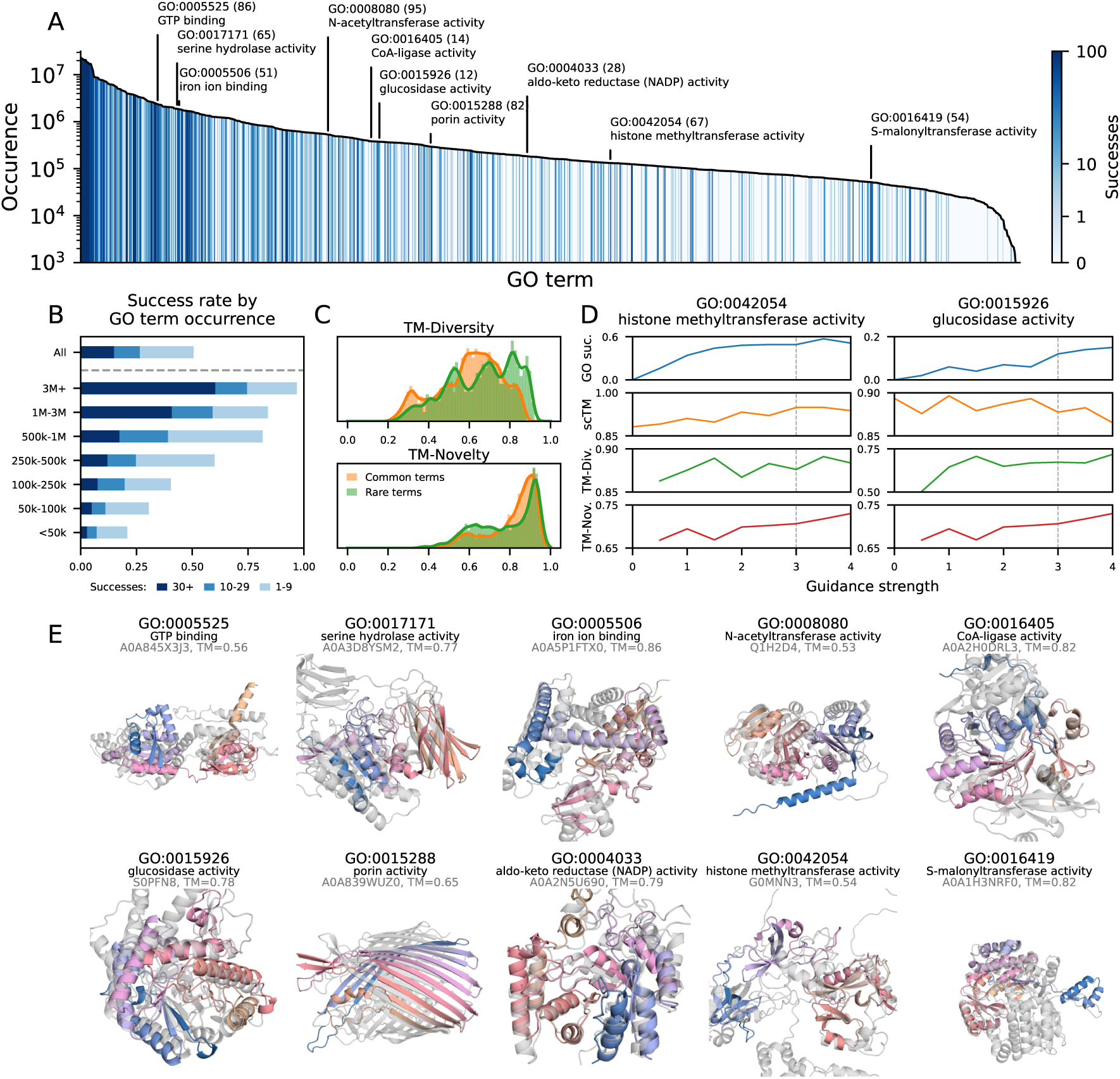
*In silico* validation of molecular function conditioning. **(A)** For each of 915 molecular function GO terms sorted by occurrence in UniProtKB, we count the number of successful generations (out of 100), where success is defined as DeepFRI probability *>* 50% for the target term. Selected terms are labeled with the number of successes. **(B)** Proportion of GO terms with different numbers of successful designs, stratified by frequency of GO term occurrence. In total, 465 terms have at least one success. **(C)** Distribution of TM-score diversity (between successful designs with the same GO term conditioning) and novelty (between designs and the most similar AlphaFoldDB entry with the same GO term) for common and rare GO terms, using a frequency cutoff of 500k. Novelty is computed only for a subset of 169 GO terms. **(D)** Impact of classifier-free guidance (CFG) for two selected GO terms with moderate success rate. For each guidance strength setting, 100 samples are generated and the success rate, mean self-consistency TM-score (scTM), diversity, and novelty are computed. Main evaluations use a guidance strength of 3. **(E)** For selected GO terms, the most novel generation is identified and superimposed onto its closest match with the same GO term from AFDB (grey).

To improve responsiveness to functional class conditioning, we explored classifier free guidance (CFG), a technique commonly used to enhance class adherence in image generative models [19]. We implement CFG with ProDiT and evaluate its ability to improve functional class conditional generation. The amount of guidance is controlled by the guidance strength parameter with unity corresponding to out-of-the-box conditional generation. We find that for GO terms with low success rates, applying CFG with guidance strengths greater than one can improve the success rate and the sequence-structure self consistency (scTM), though with some tradeoff in diversity and novelty (Fig. 2D). We expect that other established techniques from the image diffusion literature can further improve generation for less prevalent GO terms.

Prior work in function-conditioned protein generation has often cited high structural similarity and low sequence identity to known functional proteins as a sign of generalization [30, 36]; however, this does not assess generalization across structures and instead reveals memorization of structures via co-evolutionary constraints. Although the median successful ProDiT generation also has a high TM-score with known functional proteins, identifying the *most novel* design for each GO term reveals an average TM-score of 0.64 (across 169 assessed terms) to the closest neighbor in AlphaFoldDB with the same term, preserving the overall fold class but with significant local structural changes (Tab. S1). We identify and visualize these proteins for several terms (Fig. 2E) and observe alignment of substructures, indicating a shared functional site or motif but with deviation in the scaffolds. The structures are nonetheless confidently predicted by DeepFRI to possess the target GO terms, suggesting that ProDiT has learned to generate key structural elements and/or sequence motifs that underlie the functional activity without memorizing all aspects of the global structure and corresponding protein family.

#### 2.2.2 Recovery of Active Sites at Atomic Fidelity

To further verify the fidelity of successful function-conditioned designs, we investigated whether ProDiT generations recover known catalytic residues and binding motifs for molecular functions with available annotations. To do so, we developed a structural alignment pipeline to screen all GO-conditioned generated structures for the presence of labeled active site residues from UniProtKB entries with the corresponding GO term (**Methods**). Briefly, we performed a global structural alignment between successful designs and AlphaFoldDB structures with ≥2 labeled active site residues, and filtered for designs with 100% residue identity and *<*1Å RMSD for those residues. Although this pipeline is limited by the under-annotation of native active site residues in UniProtKB (across GO terms, the median percentage of structures that are sufficiently annotated is 0.23%), we found successful hits for 45 GO terms (Fig. S10, Tab. S2). In Fig. 3, we highlight two exemplar case studies that demonstrate ProDiT’s high-fidelity generation of enzyme active sites. We superimpose the generated structures against relevant PDB structures (aligned on the active site) to highlight the positioning and orientation of key residues relative to co-crystallized cofactors and substrates.

**Figure 3:**
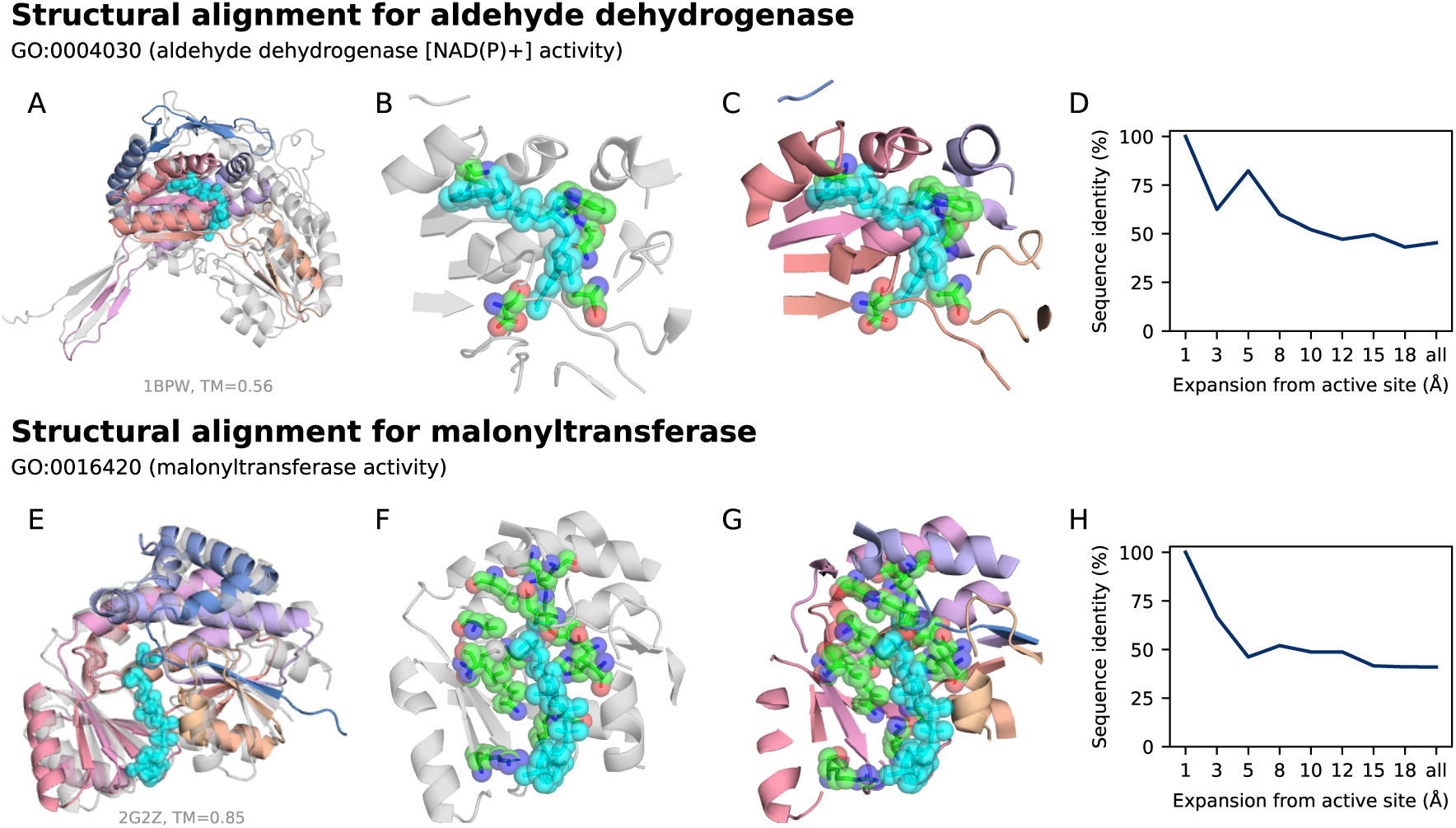
Validation of functional conditioning via structural alignment. **(A)** We superimpose a protein generated from ProDiT conditioned on aldehyde dehydrogenase [NAD(P)+] activity (GO:0004030) with PDB 1BPW, a betaine aldehyde dehydrogenase (grey), and its co-crystallized NAD+ cofactor (cyan). **(B)** View of the 10Å neighborhood of the active site in the reference protein, 1BPW. The active site residues (E263, C297) as well as key residues in the NAD+ binding motif (W165, N166, P168, K189) are highlighted. **(C)** View of the aligned 10Å neighborhood of the active site in the generated design. The active site and NAD+ binding site residues are preserved and correctly oriented. **(D)** Sequence identity versus distance from active site for 1BPW and the generated design. **(E)** We similarly superimpose a protein generation conditioned on malonyltransferase activity (GO:0016420) with PDB 2G2Z (grey), an *E. coli* malonyltransferase. The co-crystallized malonic semialdehyde and CoA ligands are also shown (cyan). **(F)** View of the 10Å neighborhood of the active site in the reference protein, 2G2Z. The active site residues (S92, H201) as well as key residues in the malonyl-CoA binding motif (R117, R190, N162, Q11, Q166, Q63, H201, H91, L194, M121, M132, S92, V168, V196, V280) are highlighted. **(G)** View of the aligned 10Å neighborhood of the active site in the generated design. The active site and malonyl-CoA binding site residues are preserved and correctly oriented. **(H)** Sequence identity versus distance from active site for 2G2Z and the generated design.

We first present a generated protein conditioned on GO:0004030, which corresponds to aldehyde dehydrogenase [NAD(P)+] activity. Aldehyde dehydrogenases catalyze the [NAD(P)+]-dependent oxidation of aldehydes to carboxylic acids and play key roles in detoxification, biosynthetic processes, antioxidant defense, and cellular regulation [3]. The generated design is aligned with the PDB structure of 1BPW—a betaine aldehyde dehydrogenase with NAD+ co-crystallized in the active site (Fig. 3A). The active site residues are generated with correct orientation, achieving an all-atom RMSD of 0.41Å (Fig. 3B,C). Furthermore, the generated protein also conserves the unique NAD+ binding motif that distinguishes aldehyde dehydrogenases from closely related alcohol dehydrogenases (e.g. Trp165, Asn166, Pro168, and Lys189 in the Rossmann fold) [24]. Strikingly, the sequence identity between the generated protein and 1BPW decreases with distance from the active site (Fig. 3D), indicating low overall homology (45% sequence identity).

We next showcase a generated protein conditioned on GO:0016420, which corresponds to malonyl-transferase activity. These enzymes play a central role in the fatty acid biosynthesis pathway by catalyzing the transfer of a malonyl group from malonyl-CoA to the acyl carrier protein. We aligned the generated design to the holo-crystal structure of a malonyltransferase from *E. coli* (PDB ID: 2G2Z), which includes malonic semialdehyde (a substrate analogue) and CoA co-crystallized in the active site (Fig. 3E). The design not only recapitulates the conserved catalytic residues (Ser92 and His201; 0.23Å RMSD) positioned to coordinate the substrate, but also preserves the topology of the malonyl-CoA binding pocket (Fig. 3F,G). Notably, residues critical for recognition of the malonyl carboxylate (Arg117), formation of the oxyanion hole (Gln11) [32] and binding of the CoA cofactor are all conserved. Nevertheless, the generated protein has only 41% sequence identity with the reference, with identical residues concentrated near the active site (Fig. 3H).

These case studies highlight ProDiT’s ability to co-generate protein sequences and structures that preserve key catalytic residues and cofactor binding sites for diverse predicted enzymatic functions. We expect that for native functions not covered by existing GO term annotations, the overall algorithm of labeling homologous sequences and tuning the model on this additional “GO-term” label should also lead to the co-generation of novel sequences and structures that preserve essential functional residues.

### 2.3 Design of Multistate Allostery

#### 2.3.1 Coupled Structure Diffusion

A significant challenge in protein design is the specification of complex, multistate functions, including allosteric regulation of protein activity [50, 29, 35, 17]. Previous efforts to design allosteric systems involve fusing domains together [35] or first exhaustively sampling alternative states using physics-based methods [17] and then searching for sequences that satisfy both states (i.e. both conformers). We hypothesized that a natively multimodal method that jointly samples sequence and structure could be well-suited for the direct generation of multistate proteins—protein sequences that can adopt multiple stable conformations. However, the scarce data for protein structural ensembles, and thus limited examples available for training, would appear to preclude any attempt to directly condition the model on properties of the target ensemble. We circumvent this hurdle by formulating *coupled structure diffusion*, a principled technique for multistate design that requires models trained only on single state sequence-structure pairs. This technique, derived from a probabilistic graphical model interpretation of the sequence-structure-function paradigm (Figure 4A), involves two structure denoising trajectories that are constrained to a single sequence denoising trajectory. We iteratively denoise the two structures using model predictions conditioned on two distinct functional motifs, while their shared sequence trajectory is constrained by the sequence of both motifs (Fig. 4B). Our protocol can be used with ProDiT out-of-the-box and requires no additional training on structural ensembles.

**Figure 4:**
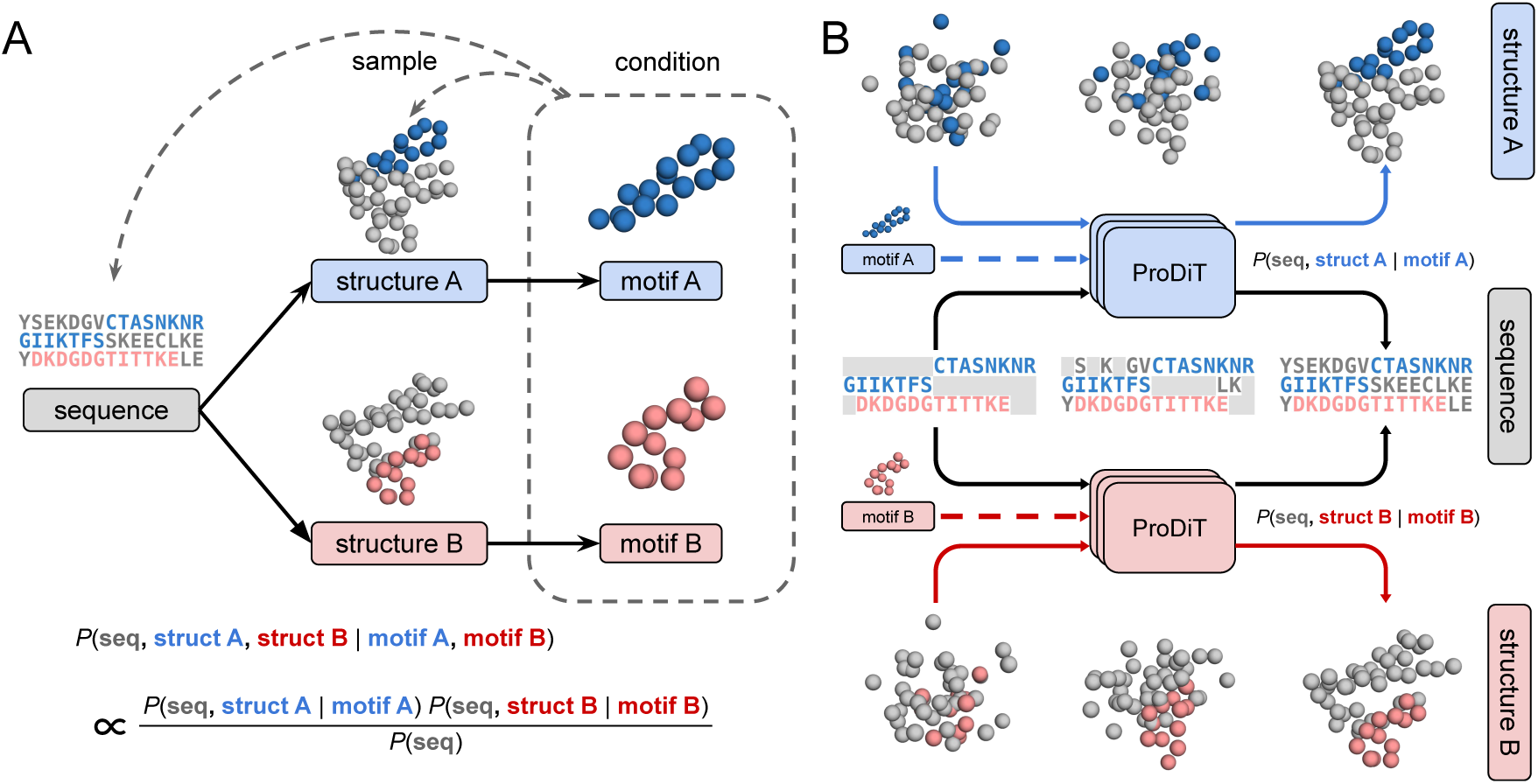
Coupled structure diffusion framework for generating multistate proteins. **(A)** The sequence-structure-function paradigm can be viewed as a directed graphical model in which structures are drawn from conditional densities *P* (struct seq) and functions (here represented as motifs) are similarly drawn from structure. Multistate proteins correspond to sampling multiple conformers from the same sequence. Hence, designing a multistate protein amounts to conditioning on the two terminal variables (motifs) and sampling the parent variables (one sequence and two structures). This conditional probability can be written as the product of conditional probabilities that only involve at most a single structure. **(B)** The product density is heuristically sampled by coupling two structure and one sequence denoising trajectories, with two evaluations of the denoising model conditioned on the two different motifs. These evaluations correspond to the two product terms in the conditional probability. A third model evaluation corresponding to *P* (seq) is not shown.

#### 2.3.2 Scaffolding Enzymatic Motifs

We explored the application of coupled structure diffusion for the scaffolding of active site motifs extracted from two enzymes, lysozyme and carbonic anhydrase, to be allosterically regulated by a calcium effector (Fig. 5). Both active sites are formed by contacts between noncontiguous segments in sequence space, which we reasoned could be disrupted by introducing an allosteric motif that induces a conformational change upon calcium binding. For each enzyme, we iteratively denoise a structure with the active site motif (taken from PDB structures 1DPX and 6LUX, respectively) and a second structure with a 12-residue EF-hand calcium binding motif from calmodulin (1PRW) randomly inserted in the sequence. Both of these structure denoising trajectories share the same sequence trajectory. To evaluate whether we obtain an active and inactive conformation, we screened the designed sequences *in silico* by predicting the protein structure with Chai-1 [42] both with and without a calcium ion. When co-folding with calcium, a substantial fraction of generated structures shifted away (in terms of RMSD) from the active site motif and towards the EF-hand motif (Fig. 5A,B,I,J). We then identified sequence designs with low active site RMSD in the unbound state, low EF-hand RMSD in the bound state, and self-consistent Chai-1 predictions for both states. This subset allowed us to interpret and analyze the designed allosteric with greater confidence. Two examples of such designs, one for lysozyme and one for carbonic anhydrase, are shown in Fig. 5

**Figure 5:**
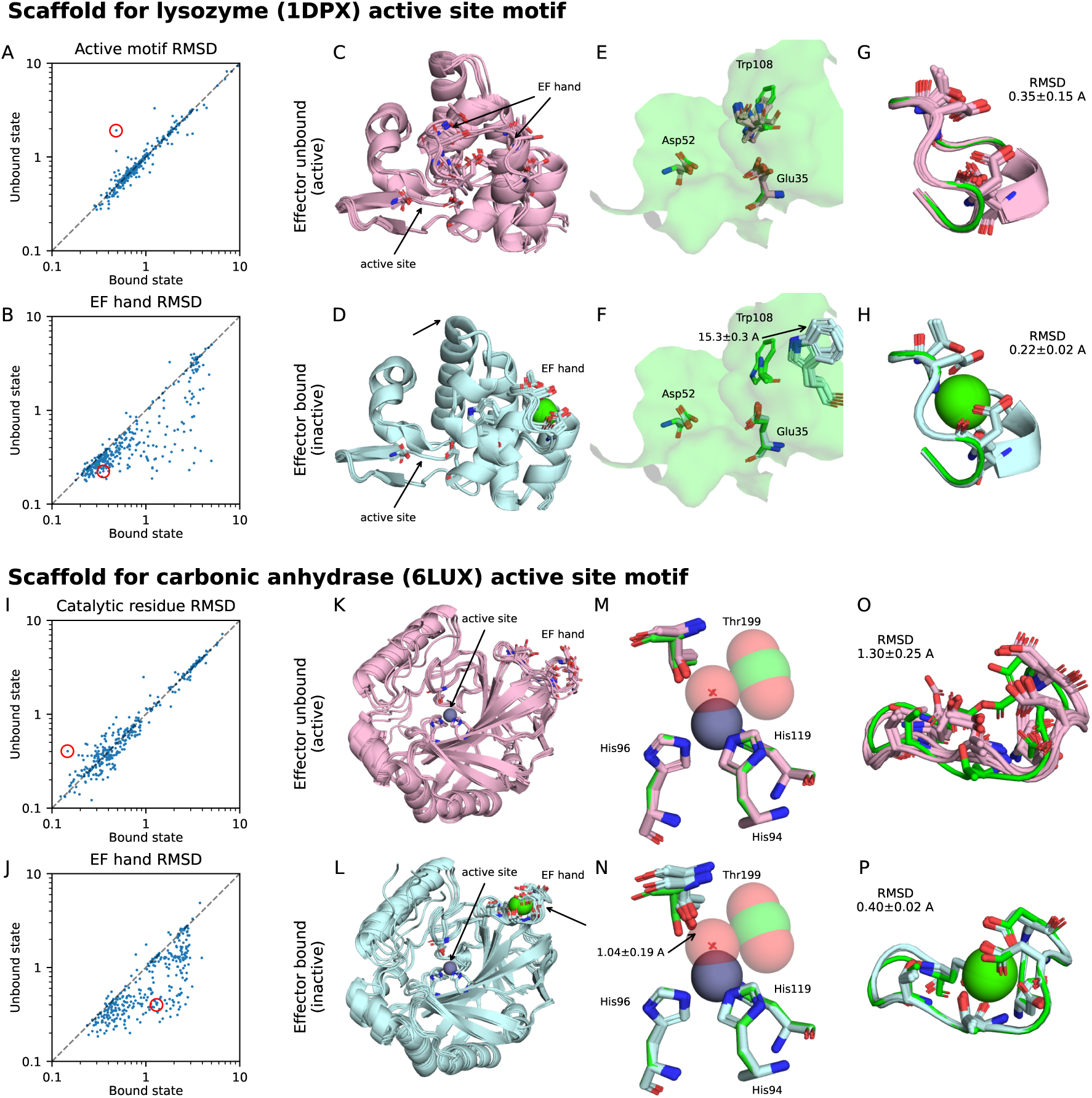
*In silico* validation of conformation switching designs. **(A)** For 320 designed scaffolds of the lysozyme motif, we refold the scaffold in both states with Chai-1 (inducing the bound state by co-folding with a calcium ion) and plot the resulting C*α* RMSDs for the four catalytic residues. The selected design is circled. **(B)** Corresponding C*α* RMSDs for the EF hand calcium binding motif. **(C)** Superimposed Chai-1 structure predictions (five total) of the unbound state. Note that the EF hand motif is not confidently predicted and adopts two conformations in the ensemble of predictions. **(D)** Superimposed Chai-1 structure predictions (five total) of the bound state. **(E,F)** The surface mesh from the native enzyme is superimposed on the native and designed motifs, highlighting the catalytic residues Glu35 and Asp52. Note the large displacement of pocket-forming residue Trp108 (full motif shown in Fig. S11,S12). We report the mean and standard deviation of the displacement of the CH2 carbon. **(G,H)** The corresponding structures of the EF-hand calcium binding sites in the unbound and bound states, with the mean and standard deviation of the C*α* RMSD calculated. **(I–P)** Similar analyses for the designed carbonic anhydrase active site scaffolds. **(M,N)** Active state geometry in the bound and unbound states, showing the carbon dioxide substrate, zinc cofactor, and nucleophilic hydroxide. Note the change in orientation of the catalytic residue Thr199 (full motif shown in Fig S13,S14). We report the mean and standard deviation of the displacement of the hydroxyl oxygen. Native motif colored in green in all subfigures.

##### Lysozyme motif

Lysozymes are enzymes that hydrolyze glycosidic bonds in peptidoglycans (EC 3.2.1.17) and are notable for being the first enzymes with a resolved structure [9]. The catalytic mechanism involves the formation of a covalent intermediate with Asp52 and uses Glu35 as a general acid to facilitate hydrolysis of a glycosidic bond [46]. This mechanism is representative of canonical glycosidase hydrolase chemistry, with the substrate specificity dictated by the surrounding microenvironment [46, 39]. For lysozymes, the Trp108 residue has been experimentally demonstrated to be critical for substrate recognition [23]. Furthermore, it is well established that carbohydrate C–H bonds interact preferentially with aromatic residues with tryptophan having a 9-fold enrichment in carbohydrate-binding modules (CBMs) [21]. In our selected lysozyme scaffold (Fig. 5A–H), Trp108 is scaffolded at its native position in the unbound state but is displaced by 15.3 Å in the calcium-bound state due to the rearrangement of several loops (Fig. 5E,F). This structural rearrangement indicates significant allosteric disruption of the substrate binding cleft, likely preventing the recognition and hydrolysis of the peptidoglycan substrate.

##### Carbonic anhydrase motif

Carbonic anhydrases catalyze the interconversion of carbon dioxide and bicarbonate (EC 4.2.1.1). This function is essential for acid-base homeostasis and is widely studied for its catalytic efficiency and potential utility in carbon capture technologies [10]. The catalytic mechanism involves a critically positioned threonine residue (Thr199) which accepts a hydrogen bond from a zinc-coordinated hydroxide ion positioned for nucleophilic attack of the substrate carbon dioxide [41, 47]. In our selected carbonic anhydrase scaffold (Fig. 5I–P), the catalytic residues are scaffolded with subatomic-level accuracy in the unbound state (0.15 Å RMSD), but calcium binding shifts the threonine hydroxyl group by ~1.04 Å in the direction of the activated hydroxide ion. This would competitively displace the Zn^2+^ coordination site occupied by the activated hydroxide nucleophile, preventing the hydration of carbon dioxide (Fig. 5M,N).

We qualify that the successful scaffolding of residues selected for the active site motif does not necessarily guarantee catalytic activity. In particular, the sensitivity of catalysis to errors in active site motif placement is unknown, and potentially important residues in the second contact shell or distal from the active site were not included in the motif-scaffolding conditioning. As such, future experimental studies will be required to verify the enzymatic activity and allosteric regulation of ProDiT’s designs. Nevertheless, our results are the first demonstration of highly controlled protein generation for allosteric disruption of enzymatic active sites, laying the groundwork for future methodological developments and fine-tuning on experimental data.

## 3 Discussion

Here, we have presented ProDiT, a multimodal protein generative model that learns a function-aware joint distribution of sequence and structure. *In silico* metrics of unconditional generation quality indicate that ProDiT matches the performance of specialized, more expensive structural models and substantially outperforms multimodal approaches that discretize structure (ESM3 [18]) or map sequences to a continuous representation (ProteinGenerator [29]). Our comprehensive benchmarking of 915 GO terms provides, for the first time, predicted functional generations across a wide range of molecular functions, often demonstrating highly accurate recovery of known active site residues in structural alignments. Enabled by ProDiT’s multimodal capabilities, we have developed a novel protocol for multistate generation, marking an important step towards the design of custom and controllable enzymes.

As researchers work to assemble a more complete picture of protein design with diverse biological functions, we anticipate ProDiT will serve as a blueprint for further developments in multimodal and multistate protein generative models. In function-conditioned design, we foresee ProDiT generations serving as new starting points for directed evolution, allowing for increased exploration of protein sequence and structure spaces. A natural extension for multistate design would be to incorporate further conditioning information relevant for protein function, such as intramolecular interactions [49] and atomic motifs [2] relevant in *de novo* enzyme design. Coupled with our unique multistate generation capabilities, such extensions could lead to particularly exciting applications in multistep enzymes, molecular motors, or signaling proteins. Ultimately, ProDiT highlights the promise of multimodal generative models to unify and advance the protein design toolkit towards better emulating the complexity of natural proteins.

## Acknowledgments

We thank Samuel Sledzieski, Alexander Shida, Arvind Pillai, Jason Yim, Woody Ahern, and Soojung Yang for helpful discussions. This work was supported by the National Institute of General Medical Sciences of the National Institutes of Health under award 1R35GM141861 (to B.B.), the NSF AI Institute for Foundations of Machine Learning (IFML), the UT-Austin Center for Generative AI, and a gift from Param Hansa Philanthropies. B.J. was supported by a Department of Energy Computational Science Graduate Fellowship under Award Number DESC0022158. A.S. was supported by a Hertz Foundation Fellowship. R.K. was supported by the Advanced Research Projects Agency for Health APECx Program and a gift from Microsoft. This research used resources of the National Energy Research Scientific Computing Center (NERSC), a Department of Energy Office of Science User Facility using NERSC awards ASCR-ERCAP0027302, ASCR-ERCAP0030607, ASCR-ERCAP0032958, and ASCR-ERCAP0027818.

## A Methods

### A.1 Multimodal Diffusion

ProDiT models protein sequence as strings in the discrete amino acid vocabulary s ∈ [1, 20]*^L^* and C*α* protein structure as an element of Euclidean space **x** ∈ R^3*L*^. The model learns to generate the data distribution by reversing noising processes defined directly on these spaces (i.e., without tokenization or embedding); as such, its training and inference formulations invoke both continuous and discrete diffusion, described below. Our training loss is a weighted combination of the structure and sequence losses, *L* = *L*_struct_ + 3*L*_seq_.

#### A.1.1 Continuous diffusion

In diffusion modeling of continuous data **x** ∈ R*^d^*, the data distribution *p*_data_ is corrupted under a forward diffusion process *d***x** = *f* (**x***, t*) *dt* + *g*(*t*) *d***w** where **w** is Brownian motion. This generates a time-evolving density *p_t_*(**x**)*, t* ∈ [0*, T*] with initial conditions *p*_0_ = *p*_data_ and *p_T_* close to Gaussian. The diffusion process can be simulated in reverse via 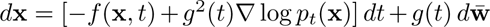 with use of the so-called score ∇ log *p_t_*(**x**) [20, 40]. We approximate ∇ log *p_t_*(**x**) with a neural network *s*_θ_(**x**) by minimizing the MSE denoising score matching objective 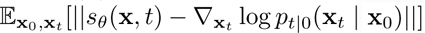 for all *t*. To apply diffusion modeling to protein structure, we take *n* = 3*L* and **x** to be the zero-mean coordinates of the C*α* atoms. We adapt, wi<u>th</u> minor notational modifications, the parameterization of Karras et al [25]: 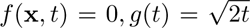 such that 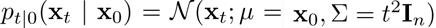. We then define time rescaling

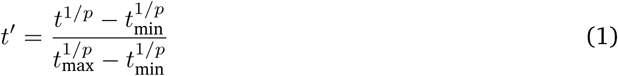

with *t*_min_ = 0.05 Å, *t*_max_ = 160 Å, *p* = 7 and train for times *t ∈*^′^ [0, 1]. That is, the neural network is trained to remove Gaussian noise ranging from *σ* = 0.05 Å to *σ* = 160 Å. The neural network score model is parameterized with post-conditioning, i.e.,

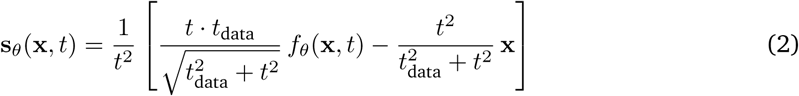

where *t*_data_ = 15 Å and *f*_θ_(**x***, t*) is the neural network. We observe that in the limit of *t →* 0 we have **s**_θ_(**x***, t*) ≍ *f*_θ_(**x***, t*)*/t* and in the limit of *t →* ∞ we have **s**_θ_(**x***, t*) ≍ (*t*_data_*f*_θ_(**x***, t*) − **x**)*/t*^2^. Hence, the network interpolates between predicting the score and predicting the clean data.

Our final structure training loss is then

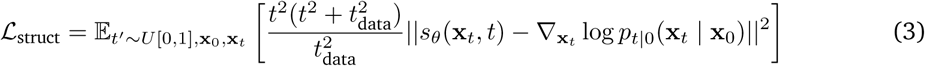

At inference time, we sample 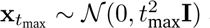 and integrate from *t*^′^ = 1 to *t*^′^ = 0 with the Euler-Maruyama scheme with uniform steps in *t*^′^. We apply low temperature annealing and modify the reverse diffusion as

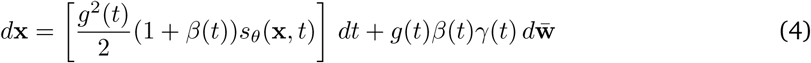

where γ(*t*) = 0.5 and *β*(*t*) = 1*/*(1 + *t/t*_data_)*^ν^*. Here *β*(*t*) controls the stochasticity via mixing level of Langevin dynamics and *γ*(*t*) controls the temperature of the dynamics. We use *ν* = 0.5 for unconditional generation and *ν* = 1 for conditional generation. As in prior work, we find these low temperature dynamics essential for generating high quality structures [22, 15].

#### A.1.2 Discrete diffusion

In diffusion modeling of discrete data [5], we represent an element of the vocabulary with its one-hot vector and transport the data distribution *p*_0_ = *p*_data_ towards a prior with probability vector ***π*** ∈ Δ*^K^*, where Δ*^K^* is the *K*-simplex. In particular, we define a corruption schedule *α_t_, t* ∈ [0, 1] such that *α*_0_ = 1*, α*_1_ = 0 monotonically decreasing and a noising process 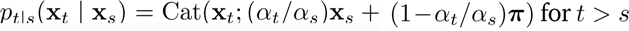. This generates noisy marginals 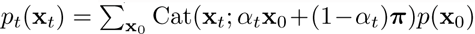. The noising process can be reversed by iteratively sampling

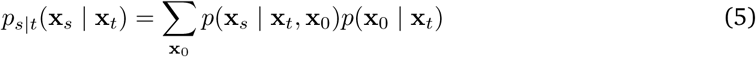

where *p*(**x***_s_* | **x***_t_,* **x**_0_) is analytically computed from Bayes’ rule and we approximate 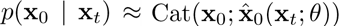 via a neural network 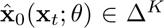. The network is trained by minimizing the cross entropy 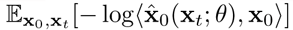 for all *t*. To apply discrete diffusion to protein sequences, we noise all positions independently and adopt the parameterization of MDLM [37] and DPLM [48]. In particular, we let *K* = 21 and define the *K*^th^ state to represent a mask state **m**. We then assign ***π*** = **m**. With this parameterization, the reverse process becomes

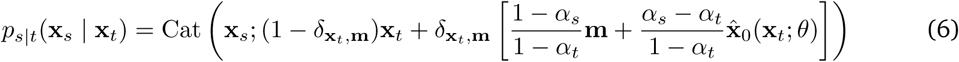

Note that the sampling is independent across positions but the neural network 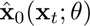 takes the entire sequence as input (for brevity we omit this distinction in notation).

Our final sequence training loss is then

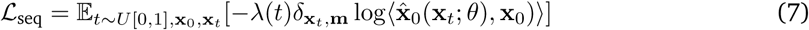

where *λ*(*t*) = 1 − *t* as in DPLM. Notably, only the logits of positions where **x***_t_* = **m** are supervised.

For inference, we initially set **x**_1_ = **m** and iteratively sample **x***_t_* with steps uniform in *t*. The sampling strategy depends on the task. For unconditional sequence generation, our sampling deviates from ancestral sampling of the reverse process *p_s_*_|_*_t_* and instead follows the approach introduced in DPLM [48], which can be cast as self-planning under a Path Planning framework [34]. Specifically, a single step from *t* to *s* with *t > s* and *L* being the sequence length proceeds as follows:

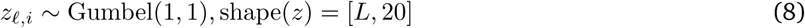

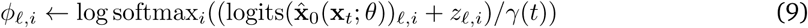

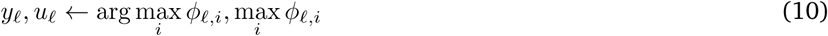

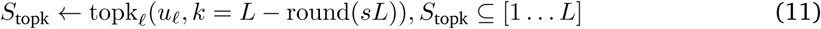

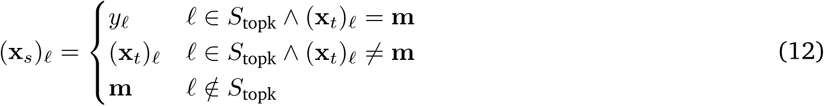

Here, *γ*(*t*) is a temperature and we set *γ*(*t*) = 0.5 for sequence generation. In co-generation, we follow a similar procedure except *γ*(*t*) = 0.1 + 0.4*t* and instead set

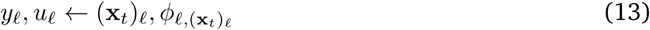

for *ℓ* such that (**x***_t_*)*_ℓ_* ≠ **m**; that is, the sampled token and log probability are replaced with the current token and its log probability under the model prediction. Finally, in inverse folding, we choose a random unmasking order and sample each position with temperature 0.1, as done in ProteinMPNN [12].

### A.2 Model Architecture and Training

ProDiT is a transformer architecture [45] based closely on the design of diffusion transformers for images [33]. In particular, it consists of 30 transformer blocks, each with a self-attention block and feed-forward layer. We adopt adaLN-Zero [33] to inject conditioning information, in particular a sinusoidal embedding of the structural diffusion time. We follow modern transformer best practices, including pre-norm, QK-norm, and GeLU activations. Unconventionally, we adopt a dropout rate of 20%, which we found essential for sequence generation performance.

The model accepts four types of inputs: structure, sequence, motifs, and GO terms. The structure is directly embedded with a linear layer without any geometric or equivariant transformations, following emerging best practice [1, 15]. The sequence is embedded with a learned embedding. The motifs are centered and also embedded with a linear layer, with motif positions additionally marked with a learned embedding in both the model input and adaLN-Zero conditioning. Finally, the GO terms are input via a learned embedding vector for each term. ProDiT then outputs via two linear heads a structural output for continuous structural diffusion and classifier logits for discrete sequence diffusion.

We train two models: one with model dimensionality 768 (without function conditioning) and one with model dimensionality 1024 (with function conditioning). These models have 321M and 576M parameters, respectively. Both models are trained with crop size 512 and batch size 250k–500k tokens across 16–32 NVIDIA H200 GPUs for approximately 500k steps. Model parameters are tracked with an exponential moving average with decay coefficient 0.999. 50% of the time, we sample motif conditioning following the motif sampling algorithm in Genie2 [27]. When GO terms are available, 85% of the time we sample a term at random in proportion to its inverse frequency across in UniProtKB, whereas 15% of the time we drop the GO term. Additionally, we completely drop the sequence or structure each 5% of the time.

### A.3 Data

Our training set is drawn from 214M proteins in UniProtKB which have associated structures in AlphaFoldDB. Unlike prior work on protein language models, we cluster proteins for training by structural similarity rather than sequence similarity; we found this to improve generation diversity. Specifically, we use the FoldSeek clusterings of AlphaFoldDB computed by Barrio-Hernandez et al [7] as hierarchical clusters; we first sample a random FoldSeek cluster, then a random MMseqs cluster within the FoldSeek cluster, and finally a random protein within the MMseqs cluster. At training time, we filter to sample proteins with pLDDT *>* 80; this results in 1.24M FoldSeek clusters, 18.9M MMseqs clusters, and 128M sequences.

To obtain function training data, we compile metadata from all UniProtKB entries and associate each entry with the set of all molecular function GO terms listed in the entry and any implied parent molecular function terms not already present (via “is_a” relationship types listed in http://purl.obolibrary.org/obo/go/go.obo). 65% of entries contain at least one non-root term, with those entries having 9.6 terms on average. We then compute frequencies of each term in our training dataset order to sample them according to inverse frequency at training time. The resulting set of 8220 terms comprises the input vocabulary of ProDiT.

### A.4 Design and Evaluation

We use the 321M-parameter model for unconditional generation protocols, as it obtains slightly better diversity, and use the 576M-parameter model for conditional generation.

#### A.4.1 Unconditional generation

For sequence generation, we sample a noisy structure at the highest noise level and fix it while iteratively unmasking the sequence. For structure generation, we initialize and fix the sequence to be the all-mask state while denoising the structure. We then design 8 sequences using ProteinMPNN, refold each of them with ESMFold, and keep the sequence with the lowest self-consistency RMSD. For co-generation, we denoise both modalities with a linear schedule and report self-consistency metrics (scRMSD, scTM) using the sequence designed by ProDiT. In all modalities, we sample generations of length *L* using *L* model forward passes. To compute the number of successful clusters, we pool all successful generations from different lengths and run FoldSeek clustering via:

~~~
foldseek easy-cluster
<DIR=
<OUT=
<TMP= --alignment-type 1 --cov-mode 0
--min-seq-id 0 --tmscore-threshold 0.5
~~~

We use the ESMFold structures for sequence generation and the raw ProDiT outputs for structure generation and co-generation.

To benchmark against prior work, we download the code and weights for ESM3 1.4B, RFDiffusion, EvoDiff 640M, ProteinGenerator from their public websites. For ESM3, because default sampling settings are not provided, we experimented with sampling hyperparameters and found the following settings to yield the best results: sequence is sampled with linear schedule, *L* steps, random order, and temperature 1; and structure is sampled with linear schedule, *L* steps, random order, and temperature 0.8. For RFDiffusion and ProteinGenerator, we use the default sampling scripts but set the number of diffusion steps to be 50 for ProteinGenerator (default 25). We use the Genie2 evaluation pipeline [27] to run ProteinMPNN on generated structures and assess all structure generation methods.

#### A.4.2 Function conditioning

The DeepFRI function prediction network outputs predictions for 942 distinct molecular function GO terms, 915 of these are in the input vocabulary of ProDiT (most of the excluded terms have been marked obselete). For each of these terms, we generate 100 proteins of length 300 by first generating a structure in 300 steps, then a corresponding sequence (using inverse folding sampling) in 300 additional steps. All sequences are refolded with ESMFold. Because some GO terms may correspond to proteins with disordered regions or flexible interdomain orientations, we do not filter based on scTM or scRMSD. The generation is marked as successful if DeepFRI predicts a probability ≥ 50% for the desired GO term from the refolded structure. We run DeepFRI with the command:

~~~
python predict.py --pdb_dir
<DIR= --ont mf --output_fn_prefix
<OUT=
~~~

To compute the diversity, we gather all successful generations with the same GO term and compute the TM-score between generations *i, j* via TMalign. The diversity of generation *i* is then the mean of *TM_i,j_, i* ≠ *j*. Note that *TM_i,j_* = *TM_j,i_* as all generations have the same length. We display histograms of diversity TM-scores of successful generations pooled across all terms (Fig. 2C), i.e., terms with more successes have higher weight.

To compute the novelty of the generations, we first compile a FoldSeek database for selected GO terms by gathering all AlphaFoldDB structures with the target GO term and running:

~~~
foldseek createdb <PDB_DIR= <OUT_DB=
We query this database via
foldseek easy-search <QUERY_PDBS= <GO_DB=
<OUT=
<TMP= --alignment-type 1
--format-output query,target,alntmscore,qtmscore
~~~

We then take the match with the highest qtmscore found by foldseek, i.e., the TM-score normalized by the length of the query (300) and assign this as the novelty TM-score of the generated protein. This ensures that high TM-scores correspond to a match for the entire generated protein rather than a short AlphaFoldDB protein matching part of the generated structure. Because of the relatively high runtime of this protocol, we only compute novelty results for 169 GO terms randomly selected from those with 10 or more success and 2M or fewer occurrences. We display histograms of novelty TM-scores of successful generations pooled across all terms (Fig. 2C), i.e., terms with more successes have higher weight.

To implement classifier-free guidance for function conditioning, we follow prior literature [19, 38] and run denoising sampling with linear combinations of logits (in discrete spaces) or scores (in continuous spaces). Let *ϕ*(**s**, **x***, t*_seq_*, t*_struct |_ ∅) and *ϕ*(**s**, **x***, t*_seq_*, t*_struct_ | *c*) represent the output of ProDiT when denoising sequence **s** and structure **x** unconditionally and conditioned on functional class *c*, respectively. Then for guidance strength *λ*, we denoise with

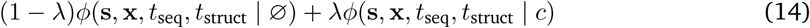

#### A.4.3 Structural alignment pipeline

To further verify that function-conditioned generations are biologically meaningful, we design a pipeline that compares the active sites of generated structures with experimental structures from AlphaFoldDB (AFDB) of the same GO term.

For each GO term, we filter UniProtKB for entries with AFDB structures and at least 2 labeled active site residues in the entry’s UniProtKB feature table (sufficiently annotated). Each generated structure is then structurally aligned to all sufficiently annotated AFDB structures with the same GO term using gemmi.calculate_superposition with selection on protein backbone atoms via gemmi.SupSelect.MainChain. For each labeled active site residue in the AFDB structure’s UniProtKB feature table, we identify the nearest residue in the generated structure with (i) the same amino acid type (ii) minimized Euclidean distance between any atom in the generated and AFDB residues. If a one-to-one correspondence was found for all annotated active site residues, we computed the all atom RMSD of the active sites. To compute the RMSD, atom positions were directly paired based on name correspondence, and the final RMSD was reported as the root mean square of the interatomic distances across the full set of matched residue pairs. Alignments with this RMSD below 1.0 Å were considered successful confirmations of active site conservation. Due to long runtimes, this process was applied to 337 GO terms.

For each successful generated structure with confirmed active site matches, we further analyzed the structure by comparing them to their five closest AFDB structures (ranked by lowest active site RMSD). For each of these structure pairs, we computed several metrics, including the global backbone RMSD, the RMSD of the active site, and the RMSDs of expanded regions around the active site using spherical expansions from 1 Å to 18 Å. All RMSD values were computed in PyMOL after realigning the structures on the selected active site or expanded region. For each level of expansion, we also match the residues in the generated structure to the closest reference residue (post-alignment) and compute the fraction of matched residue identities.

#### A.4.4 Multistate design

The objective of our multistate design protocol is to sample a single sequence **x** which folds into two distinct structures, **x**_1_, **x**_2_, supporting functional motifs **m**_1_, **m**_2_, respectively. Using the conditional independencies implied by the directed graphical model, we can rewrite this target conditional distribution as:

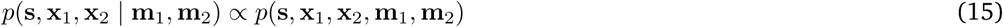

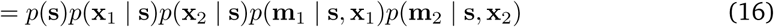

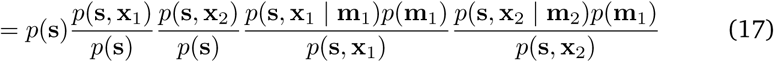

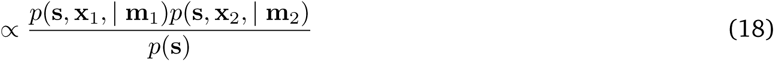

Each of these terms corresponds to our learned generative model under some form of conditioning: *p*(**s**, **x***_i_* | **m***_i_*) is the distribution sampled by motif scaffolding **m***_i_*, and *p*(**s**) is the distribution sampled by unconditional sequence generation. The product (and ratio) of these densities cannot be directly sampled by diffusion models; however, heuristic procedures for sampling similar densities involving the linear combinations of logits (in discrete spaces) and scores (in continuous spaces) are widely employed in the generative modeling literature [13], using the property that logarithms of products (and ratios) of densities are sums (and differences) of log densities.

Hence, we suggest a heuristic sampling algorithm for sampling the target density as follows: let *ϕ*(**s**, **x***_i_, t*_seq_*, t*_struct_ | **m***_i_*) and *ψ*(**s**, **x***_i_, t*_seq_*, t*_struct_, | **m***_i_*) represent the logits and score predictions, respectively, of neural networks provided noisy sequence **s**, and noisy structure **x***_i_*, diffusion times *t*_seq_*, t*_struct_, and motif **m***_i_*. Then we denoise the sequence **s** with logits

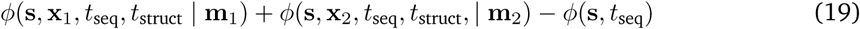

and denoise each noisy structure **x***_i_* with scores *ψ*(**s**, **x***_i_, t*_seq_*, t*_struct_ **m***_i_*). We use these expressions as logits and scores of a black-box model output and proceed with the same discrete and continuous sampling algorithms as described previously.

To explore the application this pipeline to allosteric regulation of enzymes, we posit that a multistate enzyme could be designed via two explicit structural states: one to scaffold the desired active site motif, and another to scaffold the binding motif of a desired effector. (Although there is a risk that the model would produce a degenerate design of a single or very similar structures scaffolding both motifs, we find that this does not occur in a typical sample.) We select calcium as the effector and the EF hand motif in positions 20–31 in PDB 1PRW as the effector binding motif.

For the carbonic anhydrase active site motif, we select the all residues from PDB 6LUX with C*α* coordinate within 7 Å if the position of the zinc cofactor. For the lysozyme active site motif, we first compute the midpoint of a line connecting the C*α* atoms of catalytic residues Glu35 and Asp52 in PDB 1DPX, and include all residues with C*α* coordinate within 8 Å of this midpoint. We sample 320 designs for each active site motif, using 200 (carbonic anhydrase) or 500 (lysozyme) steps of sequence-structure co-generation and motif residue indices and sequence lengths from the parent PDB. For carbonic anhydrase, we place the EF hand motif at positions 125–136 (monotonic indexing), whereas for lysozyme we randomly place the EF hand at a random non-overlapping position in each design.

To filter the designs *in silico*, we sought to identify changes that could plausibly be sufficient to disable or reduce the activity of the enzyme based on the current understanding of its catalytic mechanism. Thus, we filtered the carbonic anhydrase scaffolds based on the displacement of core catalytic residues identified in previous work. For lysozyme, we screened using the RMSD for all motif residues due to the putative importance of non-catalytic residues in substrate recognition.

### B Supplementary Results

Figs. S1, S2, S3 show additional results, broken down by protein length, on the success rates, pLDDT, and scTM of ProDiT and baselines in unconditional sequence generation, unconditional structure generation, and co-generation, respectively. Fig. S4 shows similar results for structure generation and co-generation by scRMSD.

Fig. S5 compares the scTM and scRMSD of unconditional structure generation with ProDiT when inverse folding with 8x ProteinMPNN sequences versus inverse folding with ProDiT. We note that conditional generations with GO terms use ProDiT for inverse folding. Due to degraded performance, we also explore structure generation with *ν* = 0, which produces more designable structures. We use *ν* = 0 and inverse folding with ProDiT for GO term conditioning.

Figs. S6, S7, and S8 show diversity metrics, broken down by protein length, for sequence generation, structure generation, and co-generation respectively. In particular, TM-diversity (all) pools together all pairwise TM-scores within generations of the same length, i.e., the violinplot is a density estimator over 9900 TM-scores per protein length. On the other hand, TM-diversity (max) assigns max*_i_*_̸=*j*_ *TM_i,j_* to be the diversity of generation *i*. All TM-scores are computed with TMalign.

All violin plots show the mean and inter-quartile range. Unless otherwise noted, all metrics and evaluation procedures follow their definitions in the main text.

Figs. S9 and S10 show statistics from the structural alignment pipeline for function-conditioned design.

Figs. S11 and S12 show the full lysozyme active site motif and the individual structure predictions for the selected scaffold in the bound and unbound state, along with full motif RMSDs. Figs. S13 and S14 show similar structures for the lysozyme active site motif and scaffold.

Table S1 lists the success rates, scTM, TM-diversity, and TM-novelty (when available) for all 915 GO terms evaluated for function conditioning.

Table S2 provides additional results for 45 GO terms with successful structural alignment hits.

---

**Figure S1:**
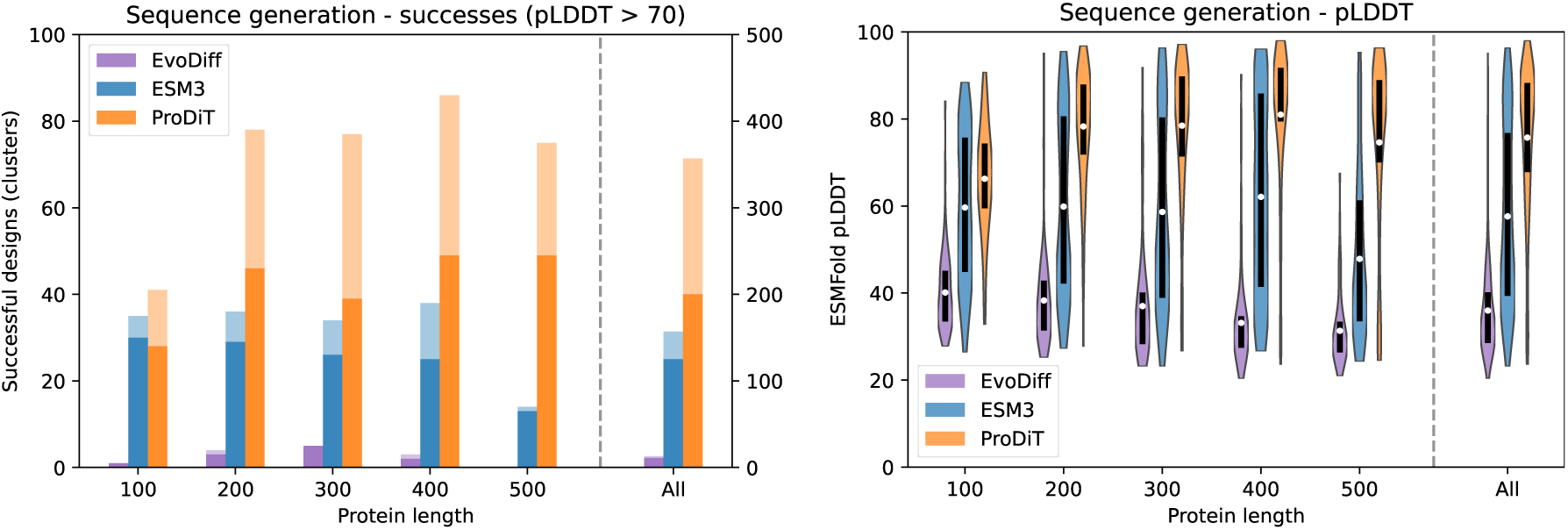
Success rates and ESMFold pLDDT for unconditional sequence generation.

**Figure S2:**
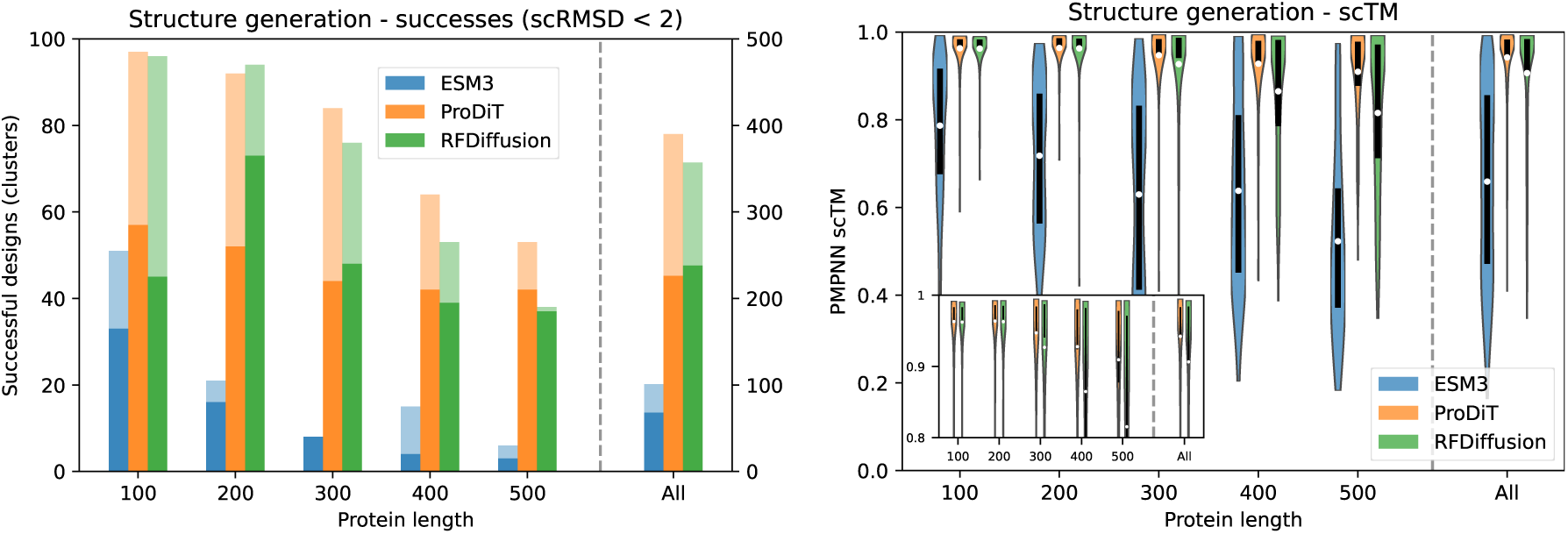
Success rates and scTM for unconditional structure generation.

**Figure S3:**
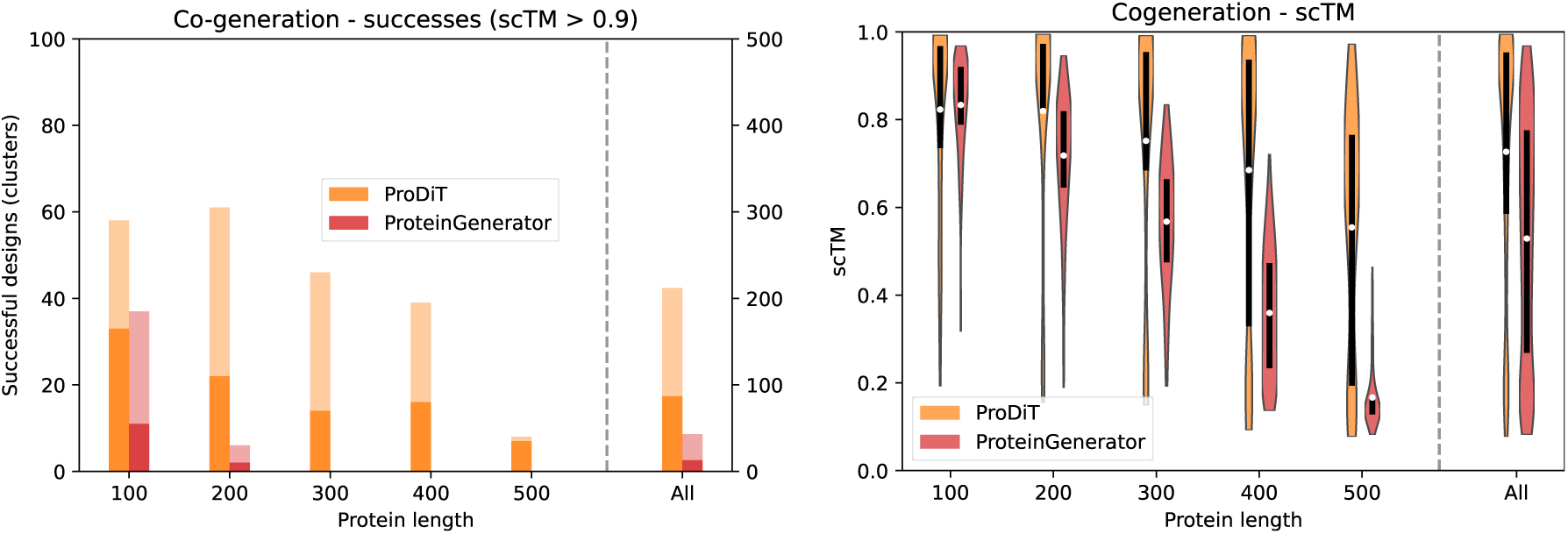
Success rates and scTM for sequence-structure co-generation.

**Figure S4:**
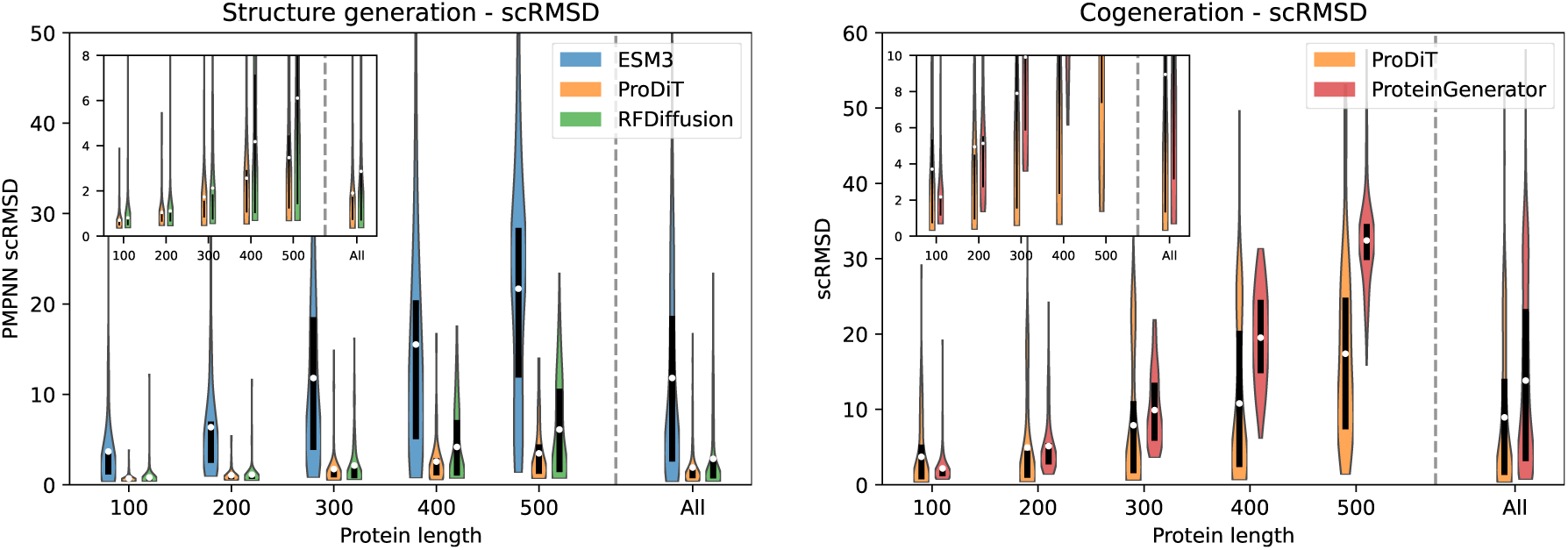
scRMSD for unconditional structure generation and sequence-structure co-generation

**Figure S5:**
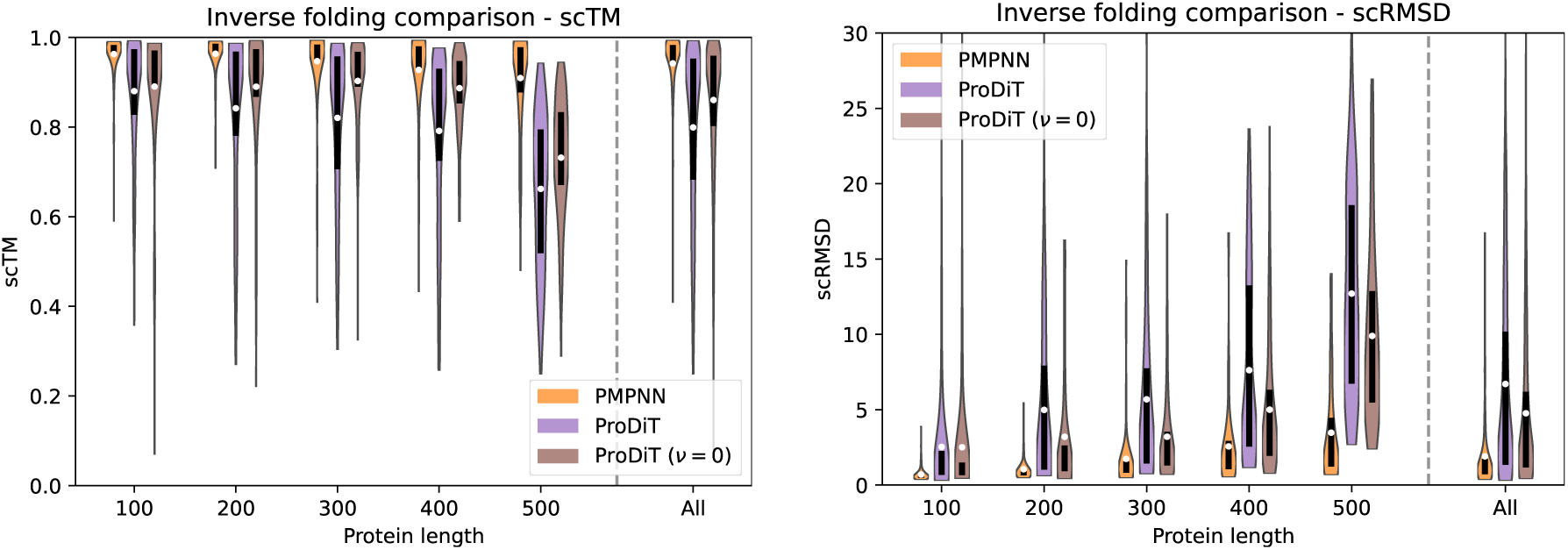
Comparison of inverse folding with ProteinMPNN vs ProDiT in structure generation.

**Figure S6:**
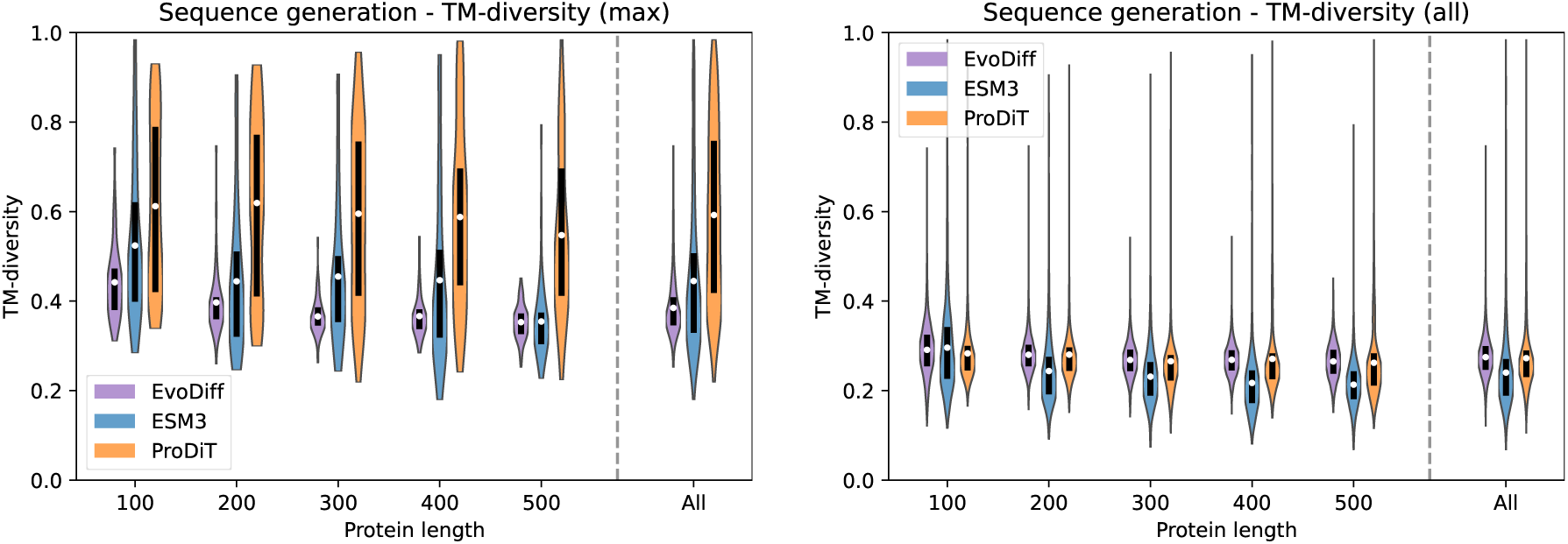
Diversity metrics for sequence generation. Lower TM-score means higher diversity.

**Figure S7:**
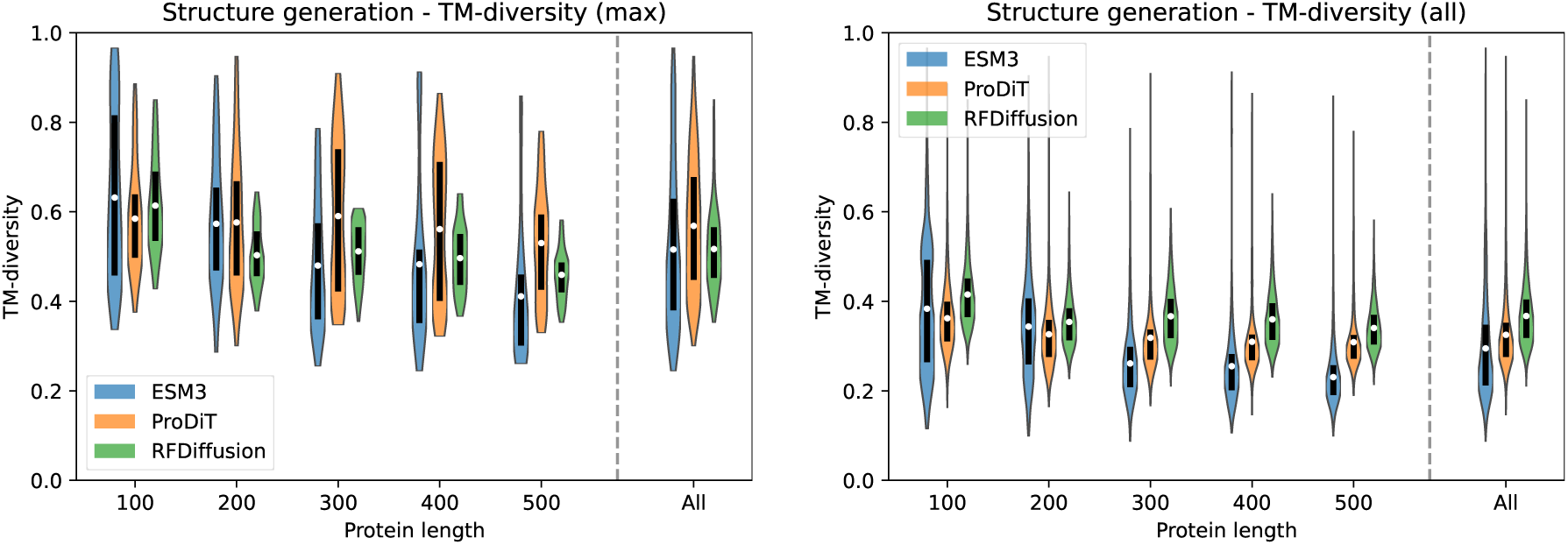
Diversity metrics for structure generation. Lower TM-score means higher diversity.

**Figure S8:**
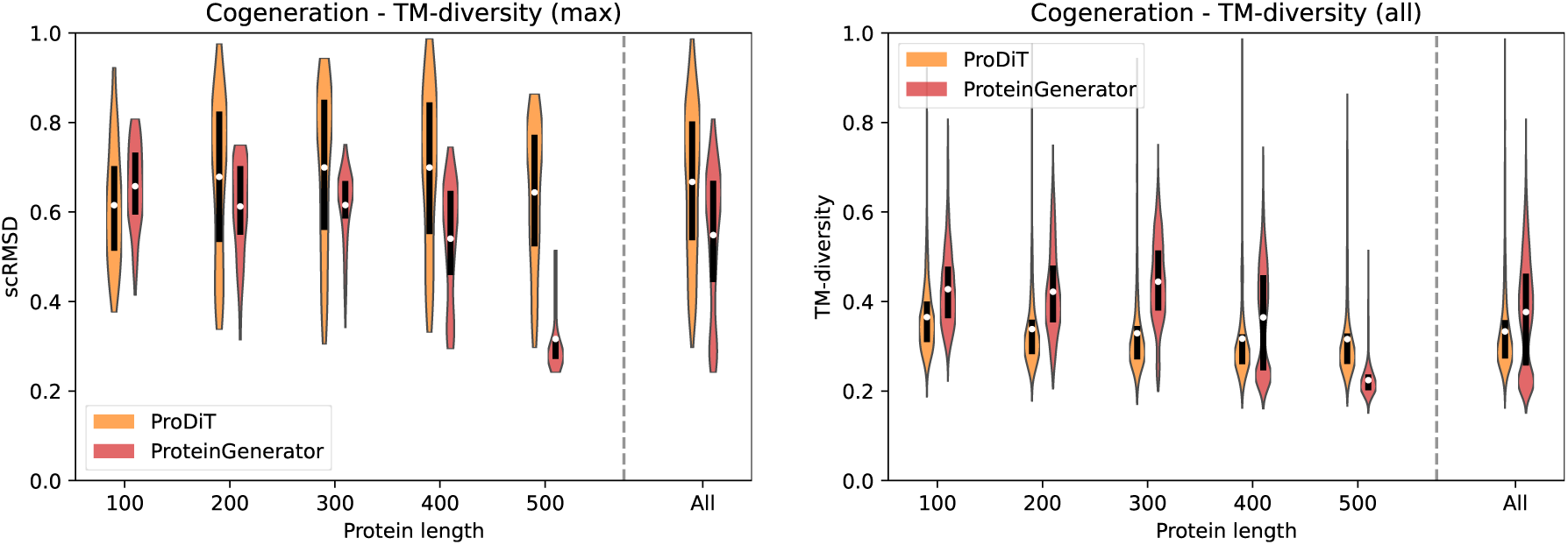
Diversity metrics for co-generation. Lower TM-score means higher diversity.

**Figure S9:**
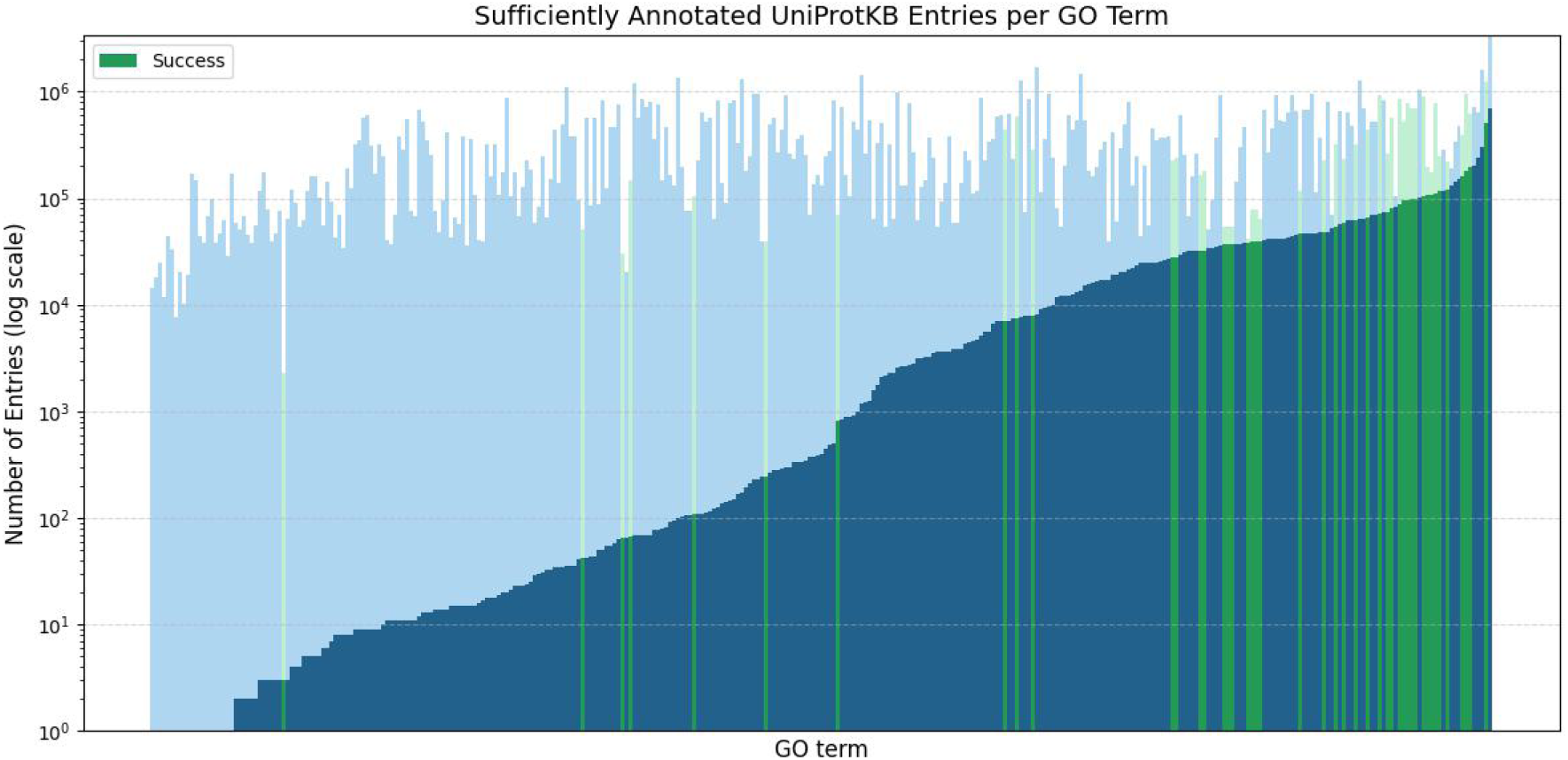
GO terms by number of sufficiently annotated UniProtKB entries. i.e., active sites with ≥2 residues. 337 GO terms are analyzed in total, out of 465 with successful ProDiT designs. Across GO terms, the median percentage of sufficiently annotated entries is 0.23%. 10 terms have zero annotated entries, and 132 terms have *<*100 annotated entries. Light bars indicate the number of total entries. GO terms with successful hits in our structural alignment pipeline (Methods) are highlighted in green.

**Figure S10:**
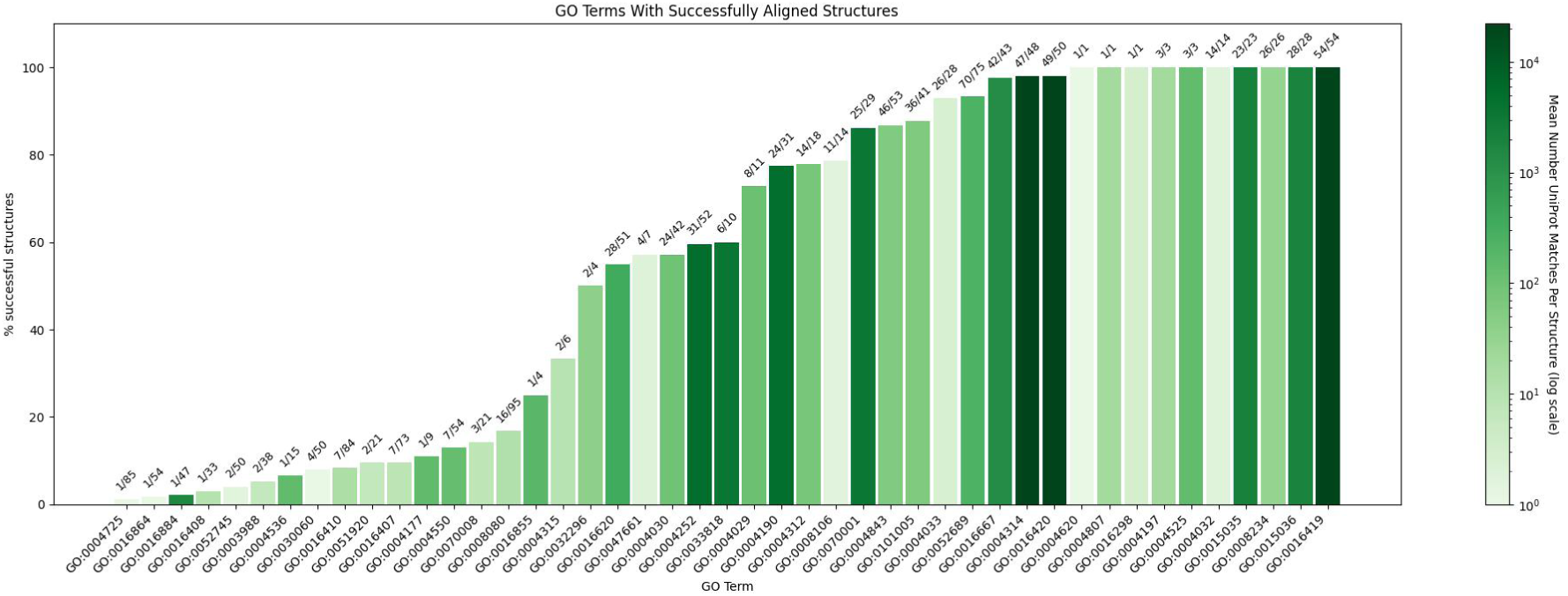
Statistics of GO terms with successful hits from the structural alignment pipeline. (45 terms). Bars are labeled with the number of successfully aligned ProDiT generations out of total successful ProDiT generations, with the bar height indicating the percentage. The bar color indicates the mean number of matching UniProtKB entries per successful design. While most GO terms matched with a few entries (median 19), a few terms exceeded 20,000 aligned hits per design.

**Figure S11:**
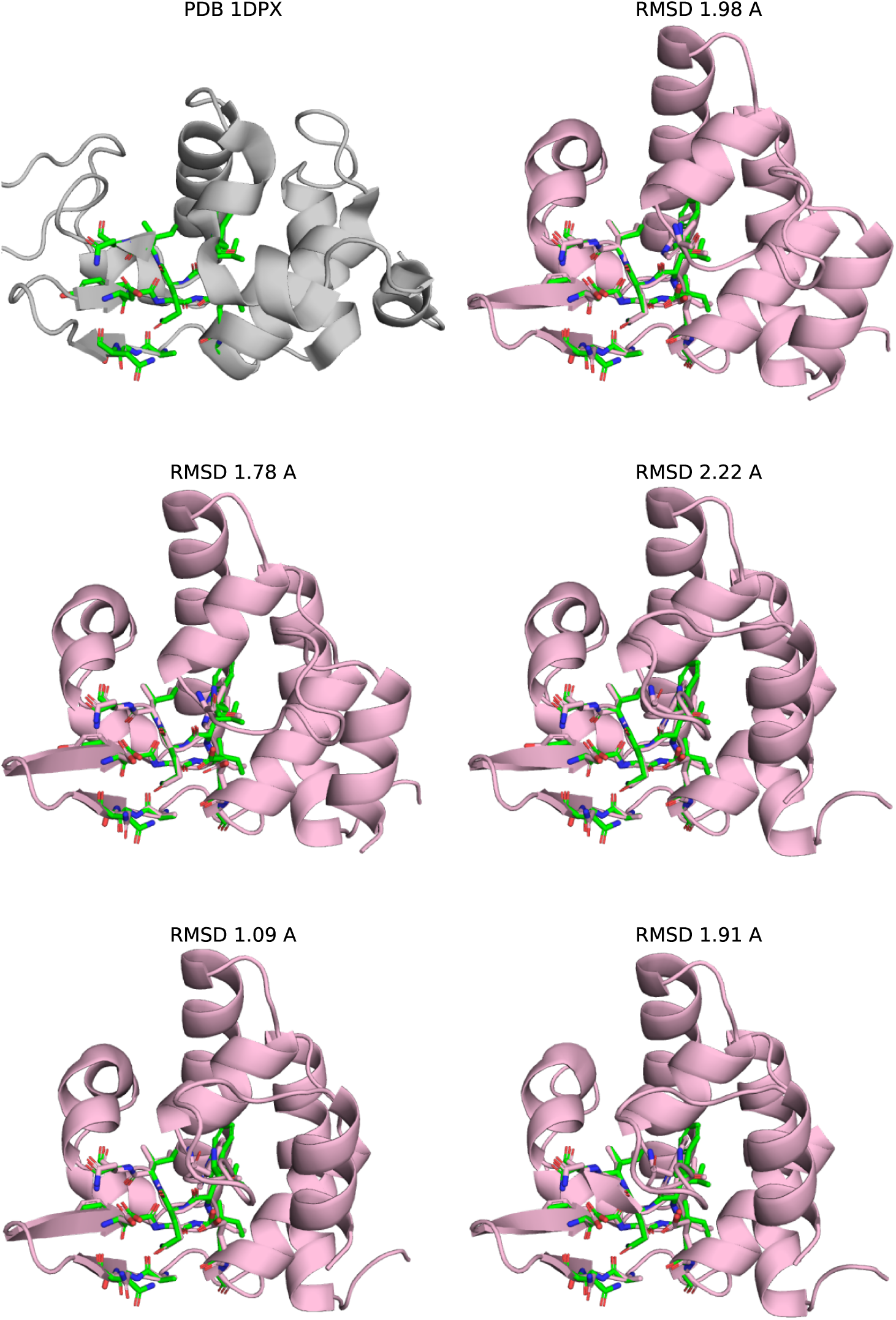
Unbound states of the lysozyme motif scaffold. Active site motif of lysozyme shown within PDB 1DPX (top left). The remaining structures show the five Chai-1 structure predictions without the calcium effector. The RMSD C*α* RMSD across all motif residues is listed.

**Figure S12:**
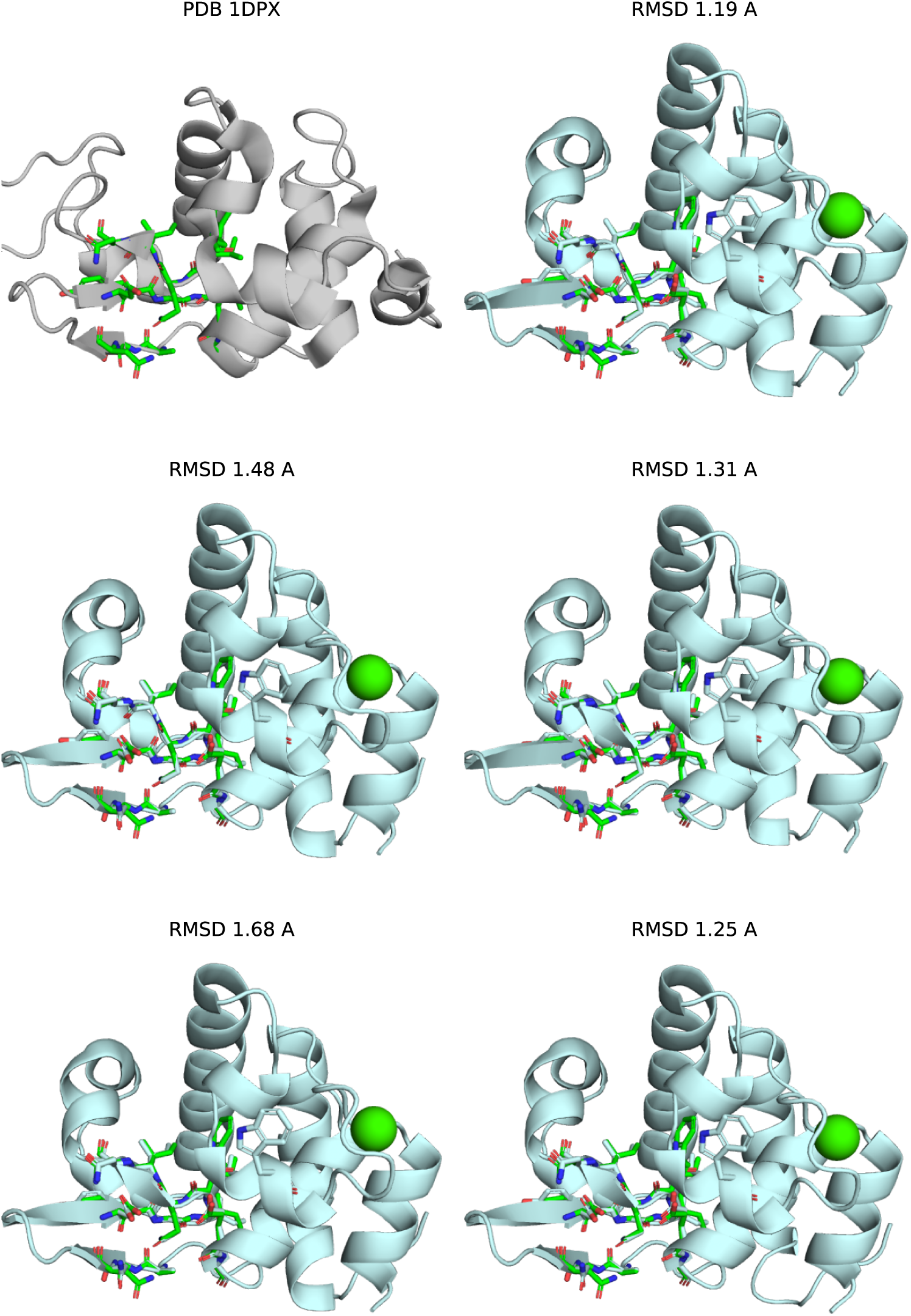
Bound states of the lysozyme motif scaffold. See previous caption.

**Figure S13:**
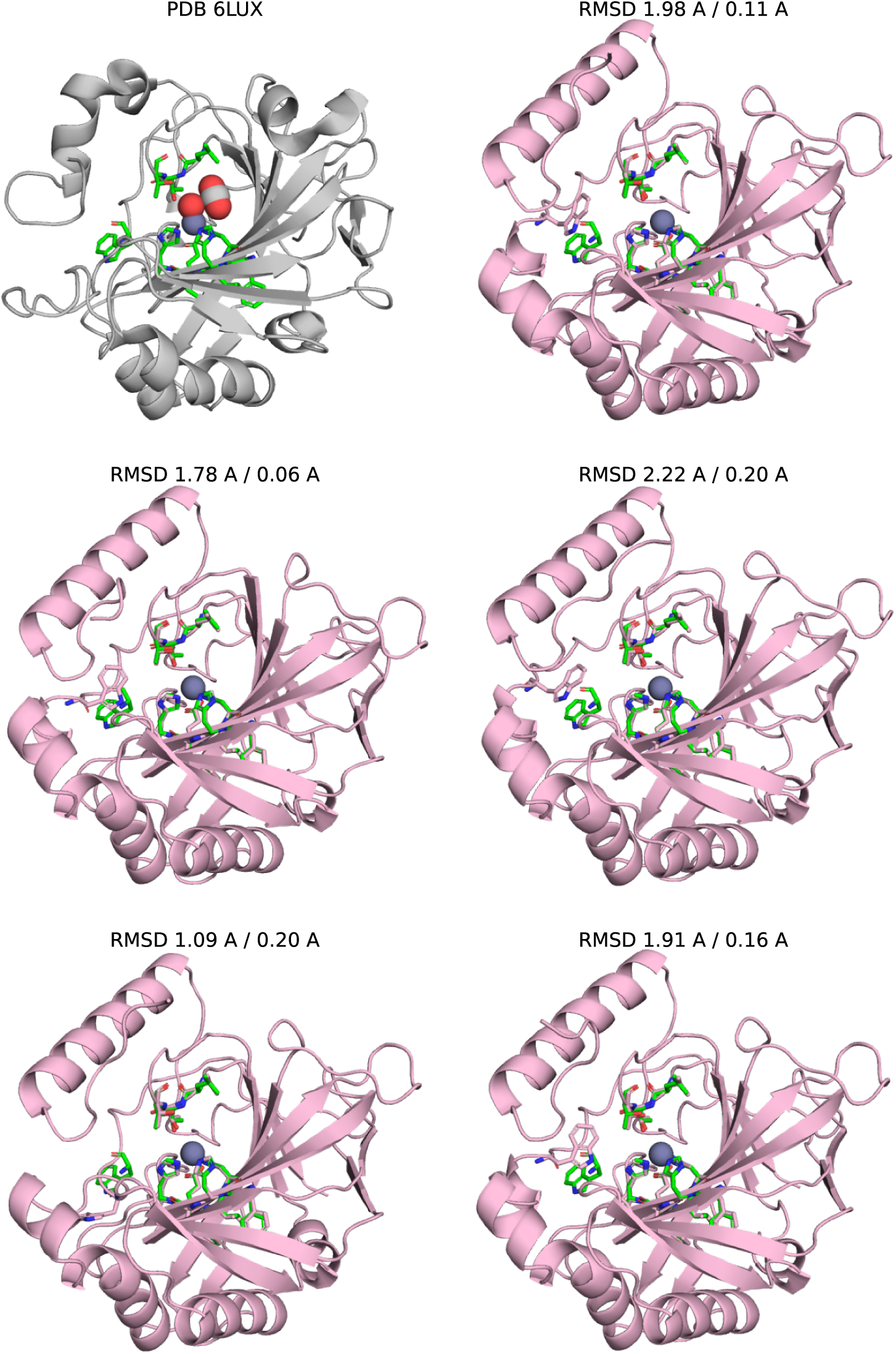
Unbound states of the carbonic anhydrase motif scaffold. Active site motif of carbonic anhydrase shown within PDB 6LUX (top left), with zinc cofactor and hydroxide and carbon dioxide substrates shown. The remaining structures show the five Chai-1 structure predictions without the calcium effector. The first RMSD listed is the C*α* RMSD across all motif residues. These are somewhat larger than typical cutoff of 1 Å and we found it was dominated by the placement of a tryptophan residue (left) whose impact on catalytic activity was unclear. Thus, we filtered based on C*α* RMSD of the four catalytic residues (second RMSD listed).

**Figure S14:**
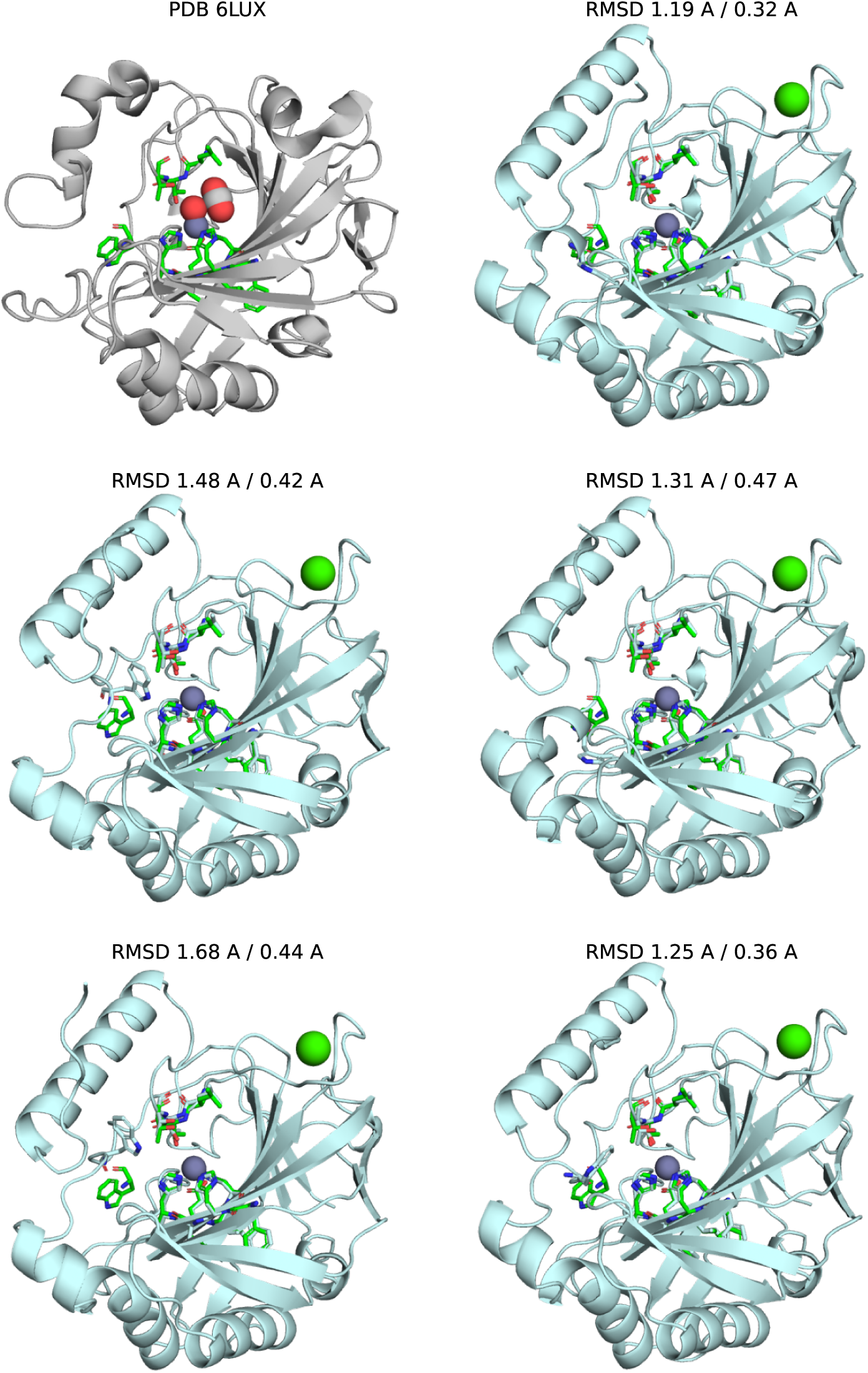
Bound states of the carbonic anhydrase motif scaffold. See previous caption.

**Table S1:**
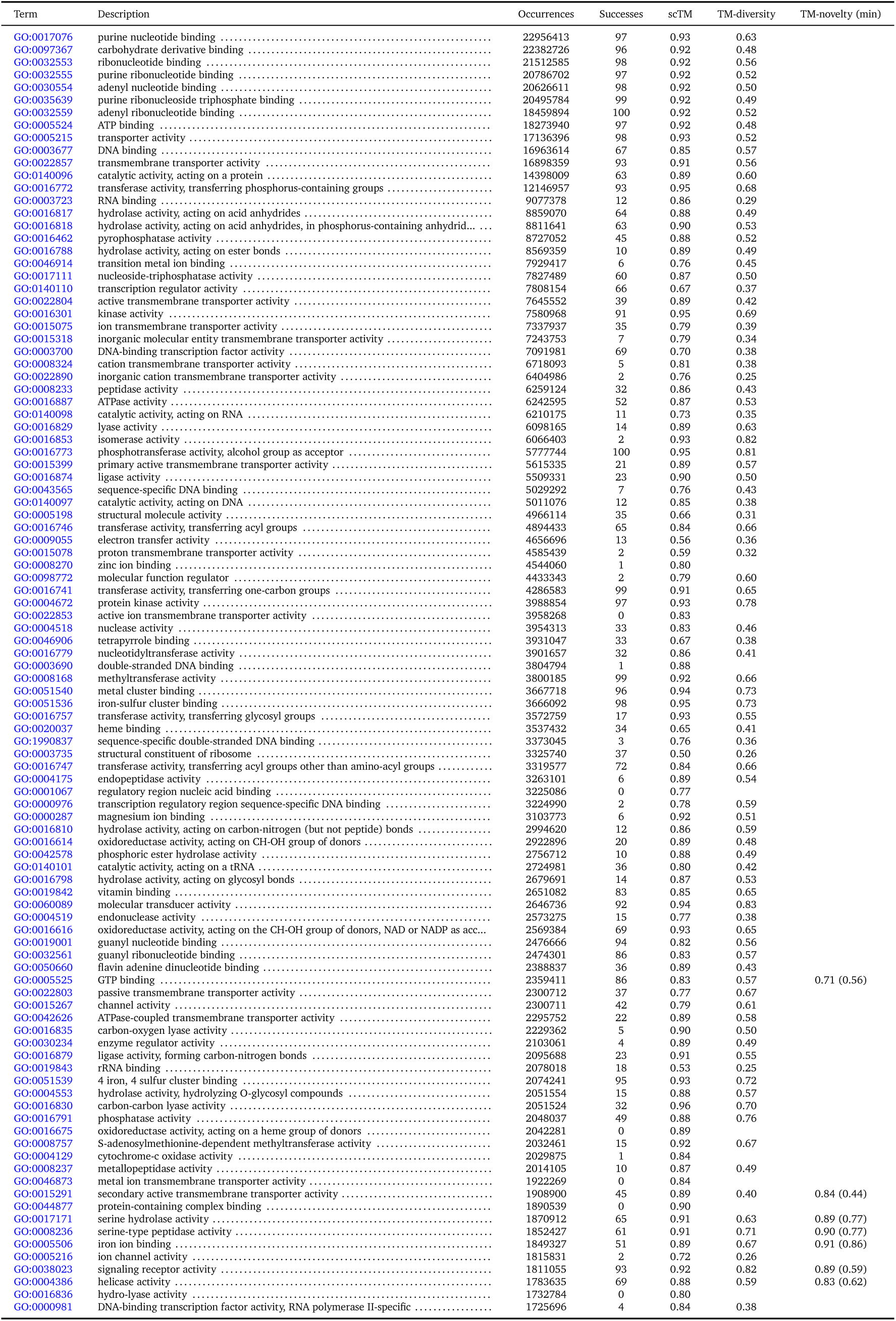
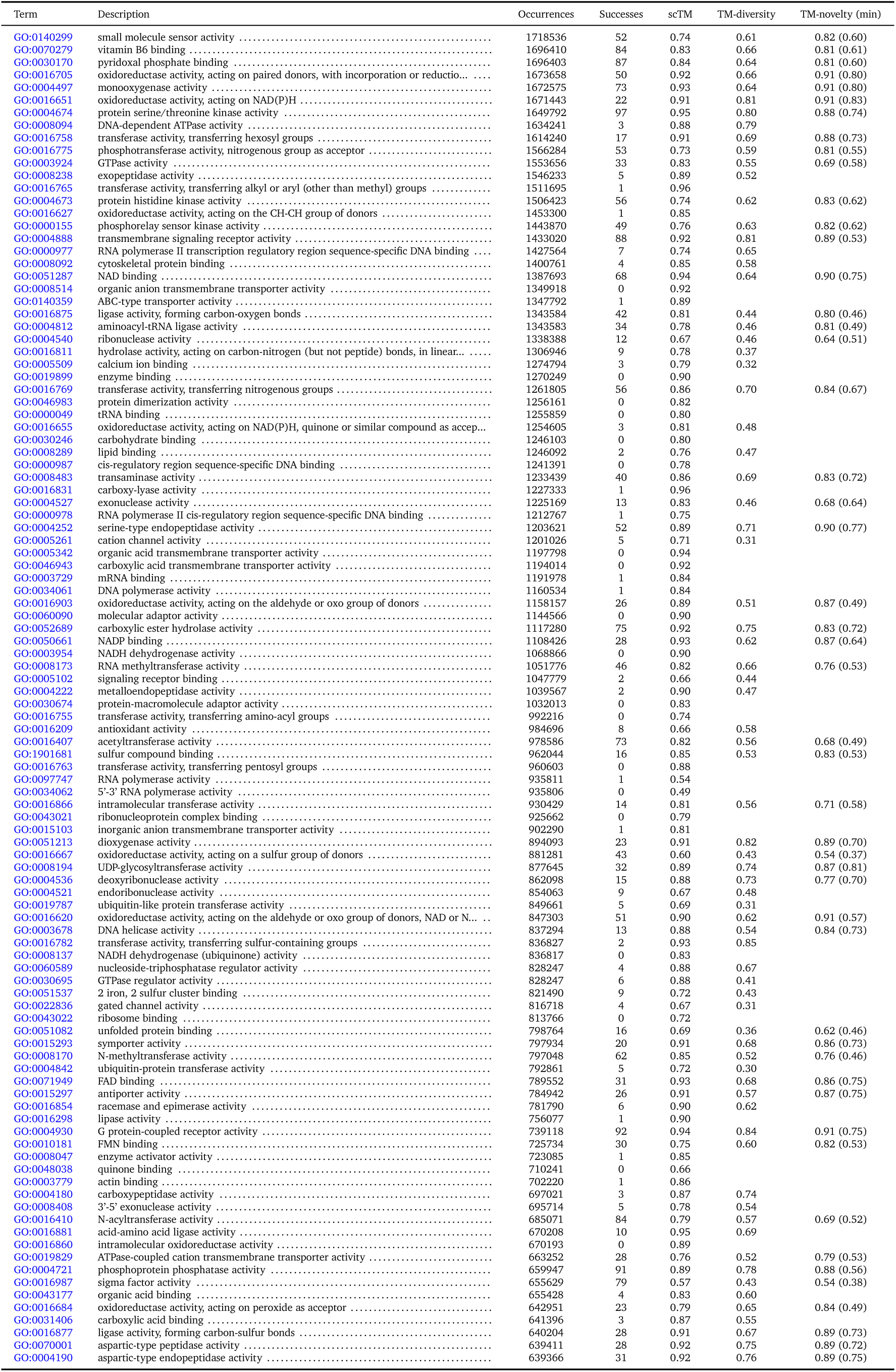
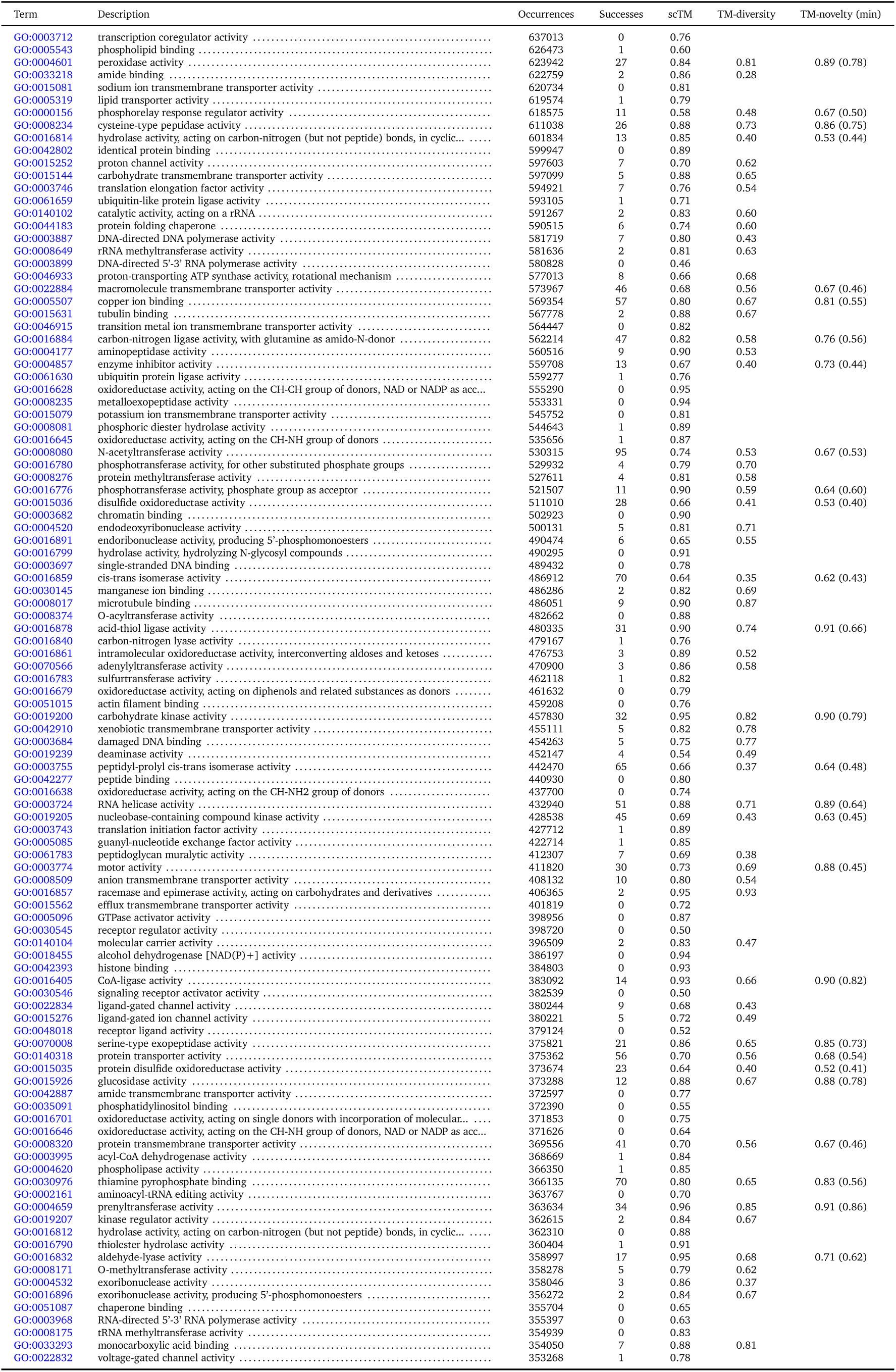
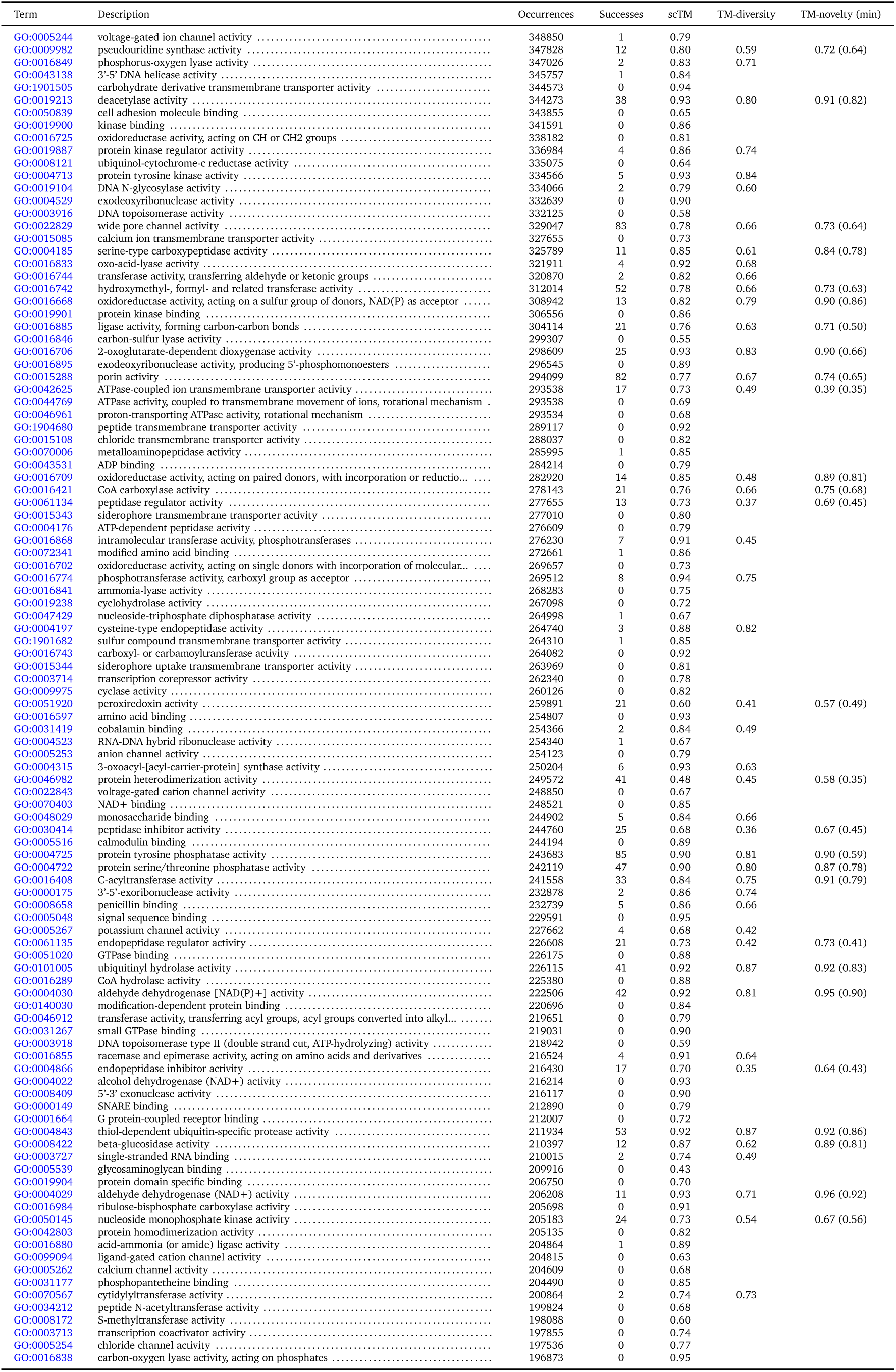
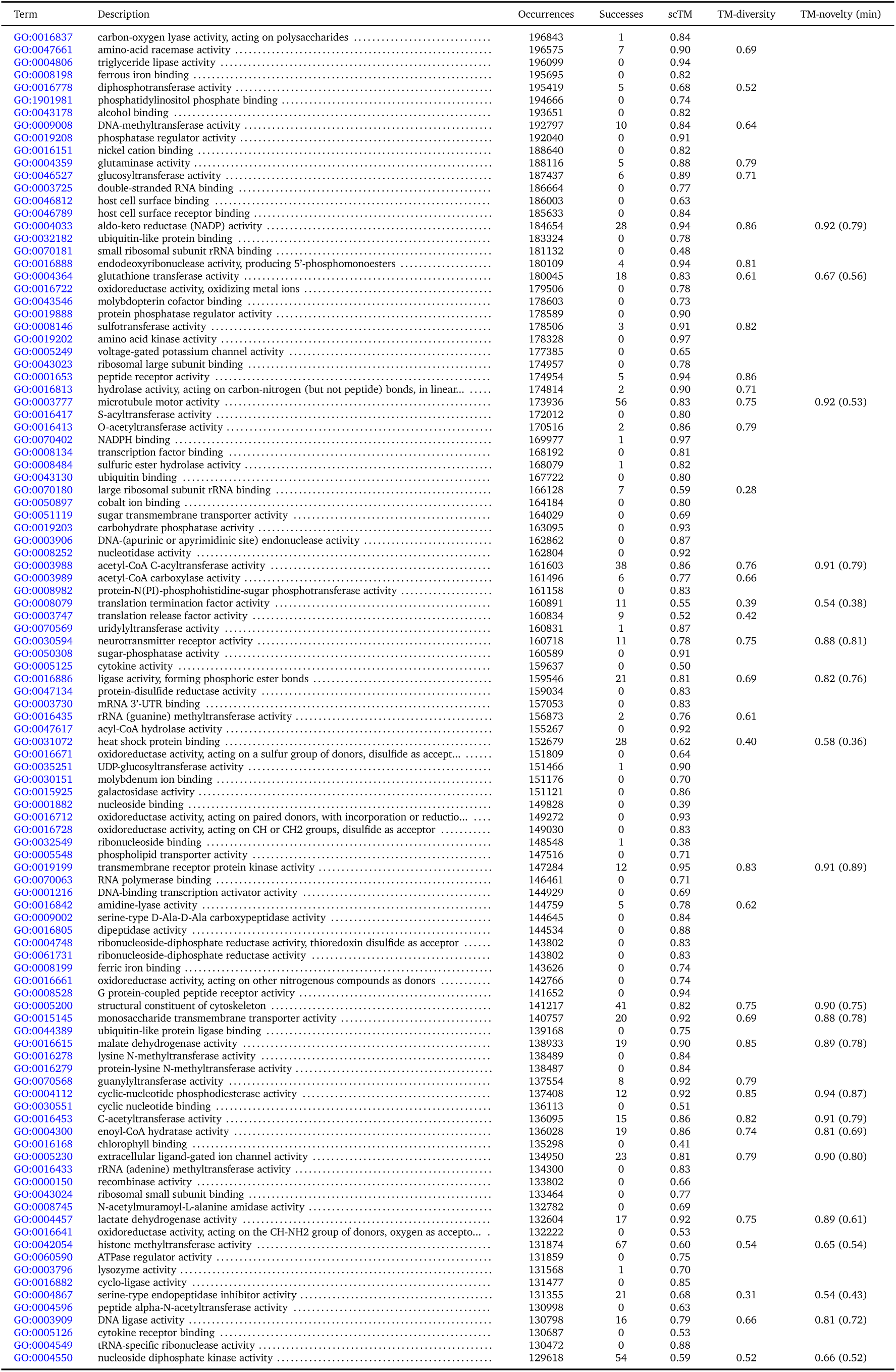
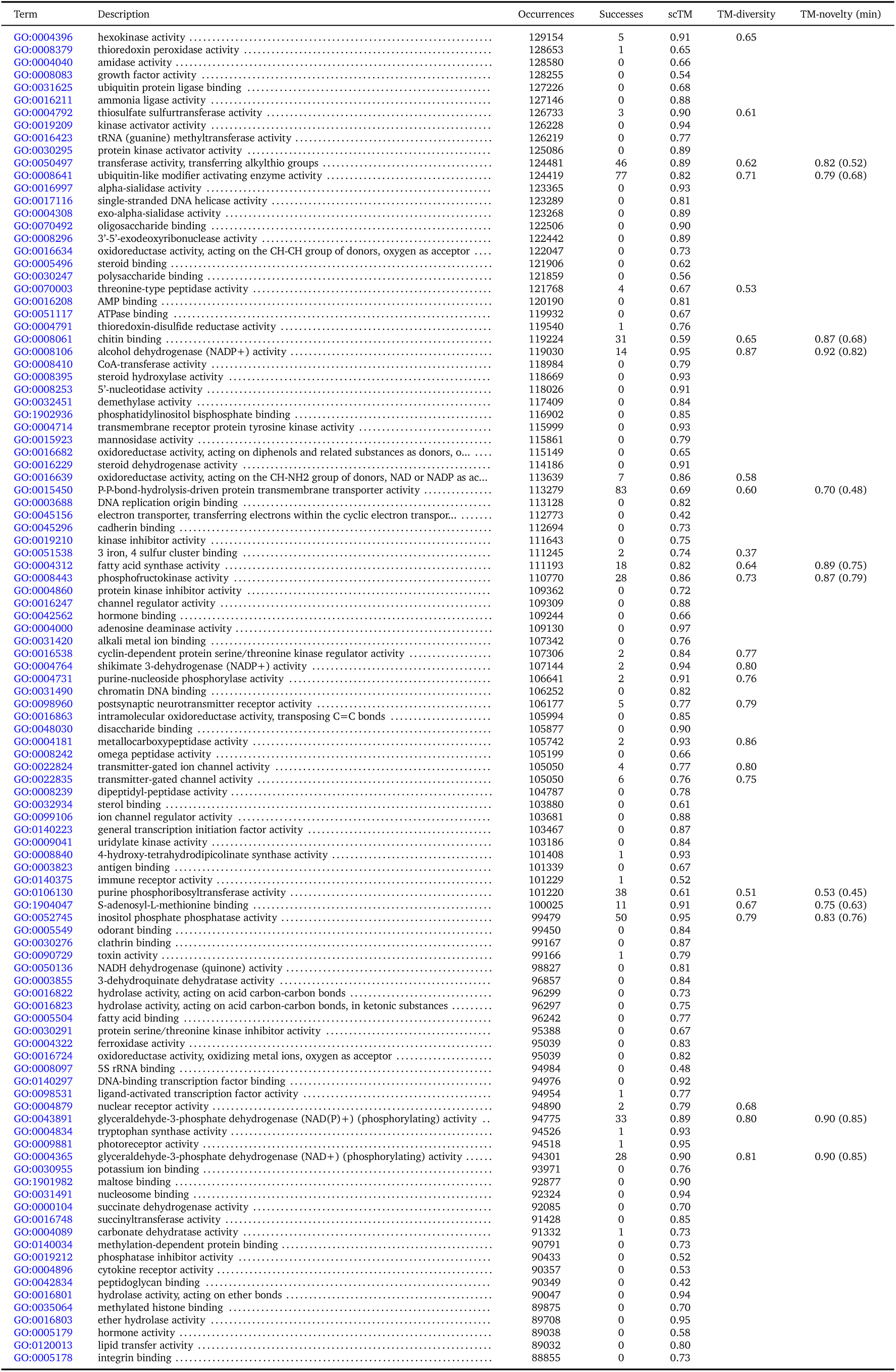
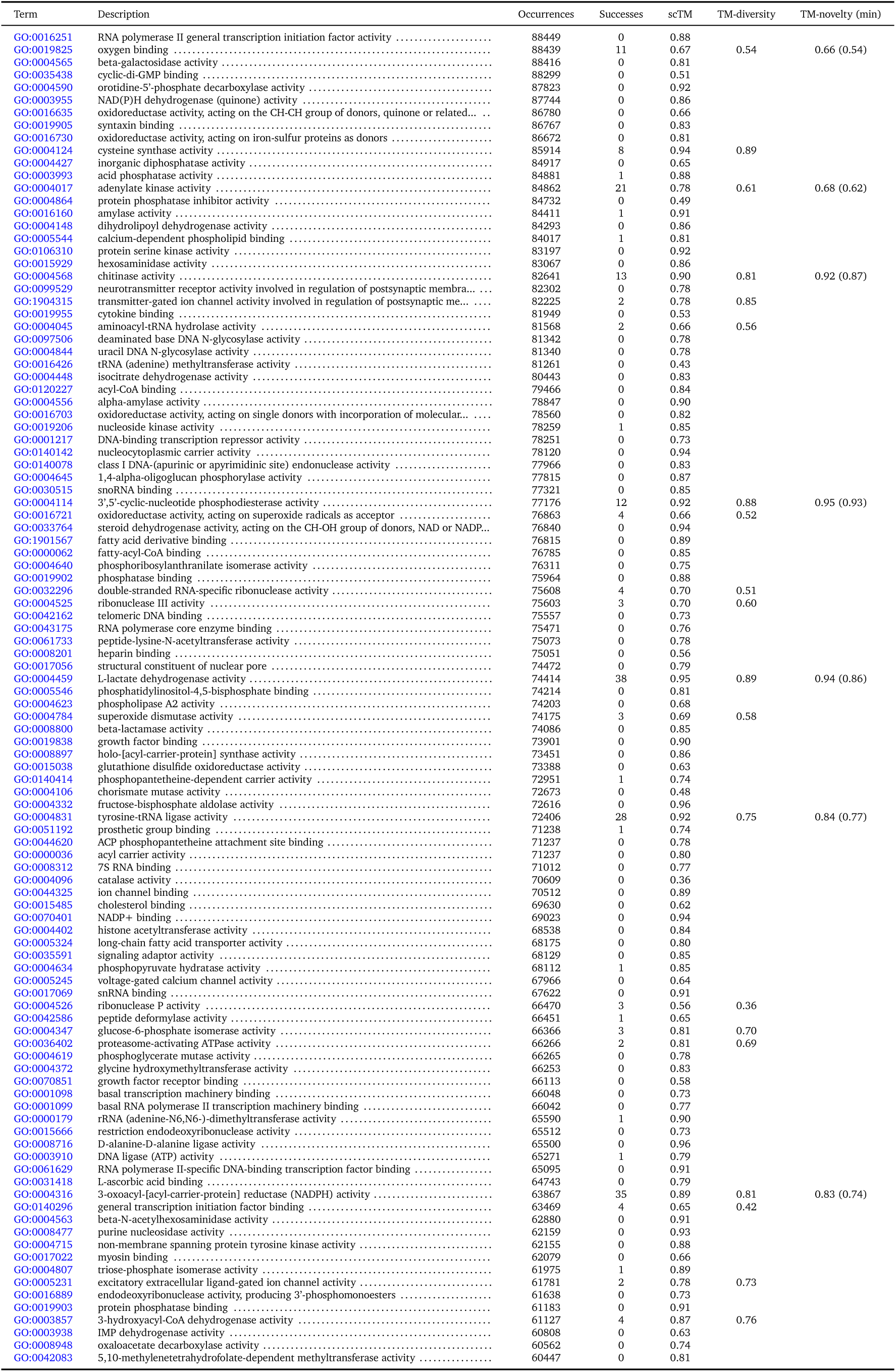
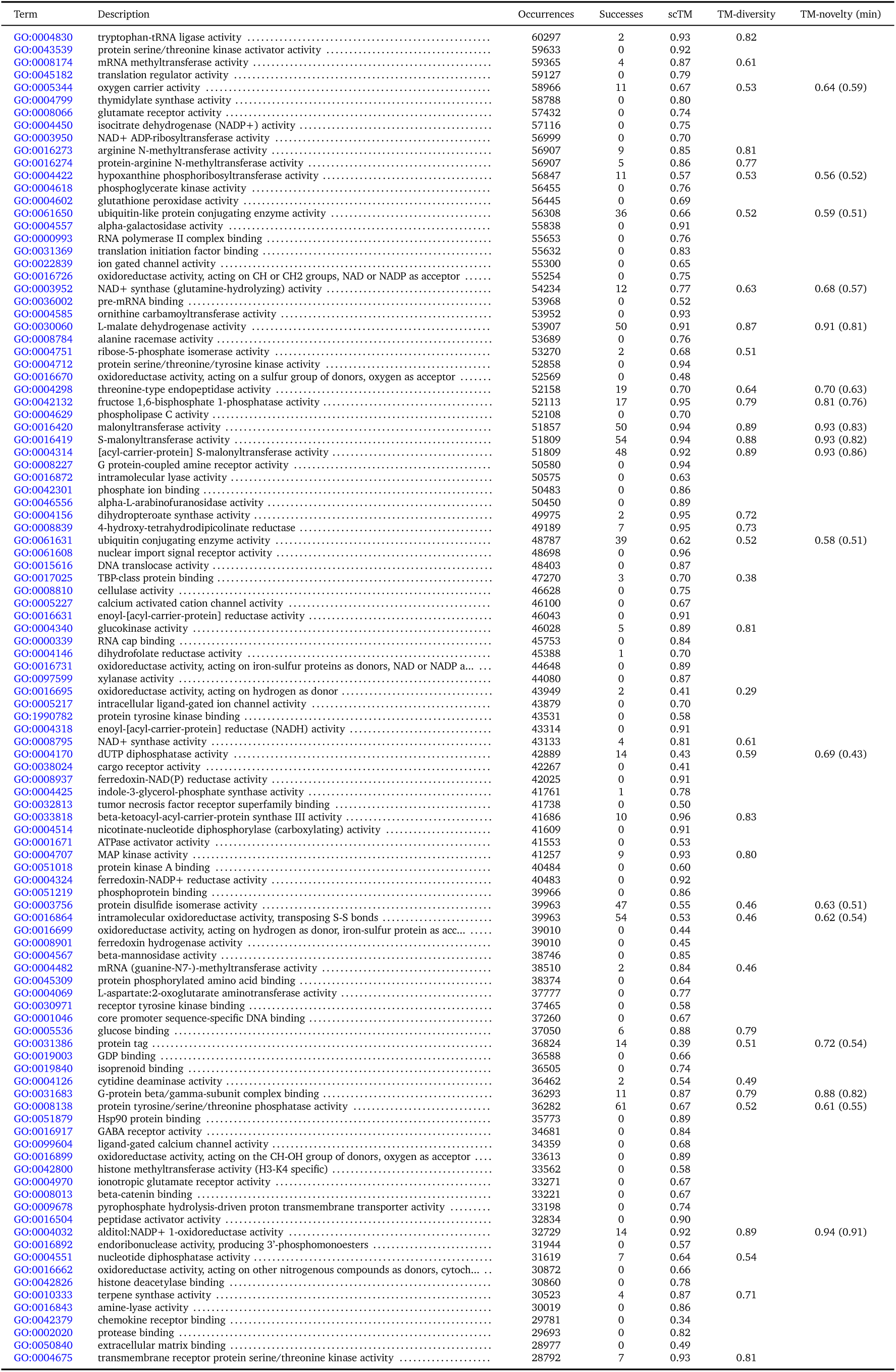
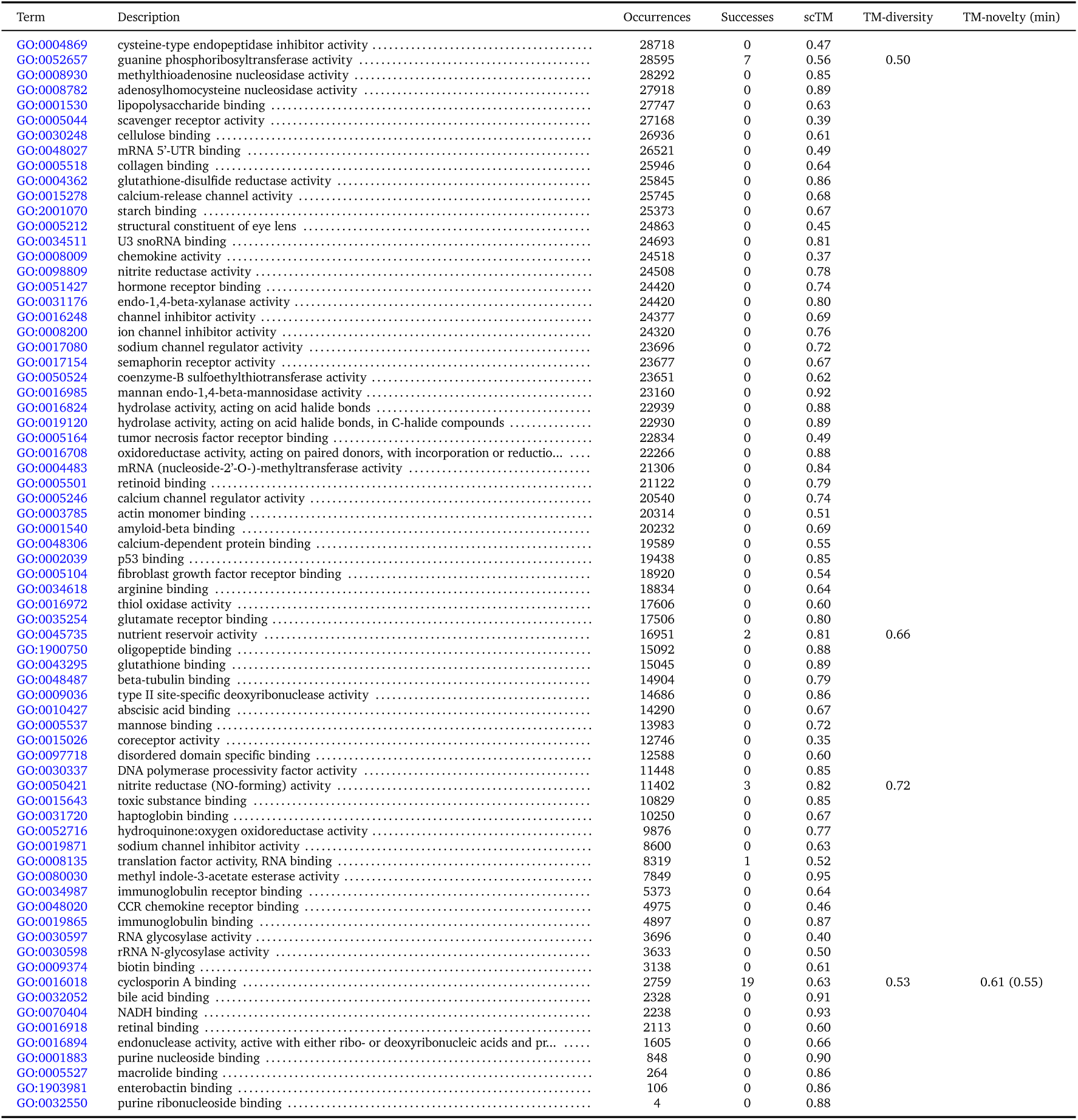
Descriptions and results for all 915 evaluated molecular function GO terms.

**Table S2:** Additional results on active site structural alignments. For each of 45 GO terms, we randomly select up to 5 successful designs and, for each design, highlight one of the aligned AFDB / UniProt structures. We report the all-atom active site RMSD, the % of matching residue identities within 5 Å of the active site, the overall sequence identity, and the active site residues annoted in UniProtKB.

| Term | Sample | AFDB / UniProt ID | Active site RMSD | 5 Å ID | Seq. ID | Active site residues |
| --- | --- | --- | --- | --- | --- | --- |
| GO:0003988 | 1 | P07871 | 0.05 | 66.67 | 48.67 | C123,C408 |
|  | 2 | Q7A2W9 | 0.16 | 63.64 | 43.0 | C86,H338 |
| GO:0004029 | 1 | A0A3Q7PKI4 | 0.13 | 80.95 | 45.33 | E210,C244 |
|  | 2 | A0A1L7TDK2 | 0.16 | 70.59 | 42.0 | E228,C262 |
|  | 3 | A0A7W3NA22 | 0.2 | 90.48 | 54.33 | E254,C288 |
|  | 4 | A0A2S5TF45 | 0.18 | 61.11 | 46.67 | E221,C255 |
|  | 5 | A0A7X3WQ03 | 0.23 | 52.63 | 44.67 | E214,C248 |
| GO:0004030 | 1 | A0A2D9E9P2 | 0.17 | 64.71 | 43.0 | E190,C224 |
|  | 2 | P52476 | 0.28 | 68.75 | 46.67 | E279,C313 |
|  | 3 | A6UQD0 | 0.24 | 66.67 | 42.33 | E240,C274 |
|  | 4 | B8M9E2 | 0.41 | 61.11 | 39.0 | E228,C262 |
|  | 5 | A0A7V9U005 | 0.15 | 72.22 | 57.33 | E260,C294 |
| GO:0004032 | 1 | O13848 | 0.28 | 57.14 | 39.44 | Y54,H109 |
|  | 2 | Q07551 | 0.13 | 45.0 | 41.67 | Y64,H122 |
|  | 3 | Q07551 | 0.25 | 54.17 | 36.0 | Y64,H122 |
|  | 4 | O13848 | 0.21 | 61.9 | 40.49 | Y54,H109 |
|  | 5 | O14088 | 0.22 | 52.17 | 36.73 | Y50,H111 |
| GO:0004033 | 1 | O14088 | 0.17 | 41.67 | 36.36 | Y50,H111 |
|  | 2 | O14088 | 0.16 | 54.55 | 39.64 | Y50,H111 |
|  | 3 | O13848 | 0.37 | 56.52 | 41.2 | Y54,H109 |
|  | 4 | Q76L36 | 0.42 | 60.87 | 41.67 | Y63,H125 |
|  | 5 | O14088 | 0.31 | 41.67 | 40.36 | Y50,H111 |
| GO:0004177 | 1 | A0A0Q4DGT6 | 0.49 | 66.67 | 51.67 | K256,R330 |
| GO:0004190 | 1 | A0A6P7MYT9 | 0.25 | 40.91 | 38.0 | I139,T325 |
|  | 2 | A0A1L9WSA4 | 0.29 | 66.67 | 43.67 | D82,D279 |
|  | 3 | A0A0D9RBZ8 | 0.29 | 72.22 | 45.0 | D97,D295 |
|  | 4 | A0A319CTT4 | 0.19 | 66.67 | 35.33 | D50,D252 |
|  | 5 | B8YJG5 | 0.16 | 65.22 | 38.0 | D108,D291 |
| GO:0004197 | 1 | Q9GL24 | 0.11 | 81.82 | 46.33 | C138,H277,N300 |
|  | 2 | Q10991 | 0.17 | 85.29 | 51.61 | C25,H163,N184 |
|  | 3 | P09648 | 0.17 | 70.59 | 49.08 | C25,H165,N185 |
| GO:0004252 | 1 | U9V2T3 | 0.15 | 58.62 | 39.06 | D18,H50,S209 |
|  | 2 | A0A7V7QN85 | 0.16 | 66.67 | 38.0 | D29,H61,S244 |
|  | 3 | A0A1I6AYN1 | 0.27 | 75.0 | 50.67 | D178,H217,S434 |
|  | 4 | A0A1I5DDV9 | 0.15 | 77.42 | 49.33 | D62,H110,S292 |
|  | 5 | A0A7X2J0A2 | 0.17 | 79.41 | 44.33 | D137,H174,S352 |
| GO:0004312 | 1 | A0A6L7X5E9 | 0.22 | 54.17 | 41.33 | S128,H233 |
|  | 2 | A0A535J0M4 | 0.46 | 45.83 | 39.0 | S103,H212 |
|  | 3 | A0A3M1LXU7 | 0.48 | 47.83 | 39.33 | S94,H201 |
|  | 4 | A0A2E6YIR2 | 0.3 | 60.0 | 37.0 | S103,H208 |
|  | 5 | A0A2M8ETE7 | 0.2 | 42.86 | 33.0 | S114,H219 |
| GO:0004314 | 1 | A0A352PU99 | 0.31 | 26.92 | 35.33 | S91,H196 |
|  | 2 | A0A496AKM2 | 0.2 | 57.69 | 38.67 | S96,H204 |
|  | 3 | A0A2J0KPU7 | 0.18 | 53.85 | 38.0 | S145,H250 |
|  | 4 | Q5F4X7 | 0.2 | 53.85 | 37.0 | S90,H199 |
|  | 5 | A0A1G3UH18 | 0.21 | 65.38 | 31.33 | S92,H202 |
| GO:0004315 | 1 | P9WQD9 | 0.63 | 37.93 | 44.67 | C171,H311,H345 |
|  | 2 | P0AAI7 | 0.24 | 56.67 | 43.0 | C164,H304,H341 |
| GO:0004525 | 1 | A0A0F2Q590 | 0.14 | 56.52 | 30.29 | D54,E126 |
|  | 2 | A0A535S225 | 0.05 | 56.52 | 34.91 | D54,E124 |
|  | 3 | A0A2D6TAR6 | 0.16 | 56.52 | 34.03 | D49,E121 |
| GO:0004536 | 1 | A0A351R9X6 | 0.31 | 51.52 | 29.69 | Y104,D145,H246 |
| GO:0004550 | 1 | A0A2J7QC69 | 0.32 | 60.71 | 39.67 | H162,H316 |
|  | 2 | A0A803K9Z7 | 0.27 | 57.14 | 40.0 | H212,H361 |
|  | 3 | D8TIX5 | 0.4 | 65.52 | 40.67 | H203,H354 |
|  | 4 | A0A2J7QC53 | 0.31 | 60.71 | 41.33 | H231,H385 |
|  | 5 | A0A2J7QC66 | 0.4 | 48.15 | 39.67 | H208,H362 |
| GO:0004620 | 1 | P16233 | 0.21 | 50.0 | 36.0 | S169,D193,H280 |
| GO:0004725 | 1 | A0A6M8FQ69 | 0.27 | 21.43 | 35.06 | C7,R13,D123 |
| GO:0004807 | 1 | A0A840WL33 | 0.26 | 81.82 | 34.57 | H93,E163 |
| GO:0004843 | 1 | Q8LAM0 | 0.31 | 28.57 | 38.67 | C32,H310 |
|  | 2 | G5E8G2 | 0.09 | 66.67 | 41.67 | C60,H307 |
|  | 3 | A0A7D8YLP6 | 0.06 | 71.43 | 50.0 | C311,H741 |
|  | 4 | A0A0J8UXP0 | 0.28 | 38.1 | 48.0 | C226,H657 |
|  | 5 | O57429 | 0.21 | 55.56 | 42.0 | C28,H309 |
| GO:0008080 | 1 | A0A2E8FZC6 | 0.13 | 48.0 | 43.31 | E109,Y121 |
|  | 2 | A0A7V7MUE6 | 0.14 | 50.0 | 48.57 | E97,Y109 |
|  | 3 | A0A2W1KFP4 | 0.13 | 53.85 | 43.04 | E103,Y115 |
|  | 4 | A0A411WMN2 | 0.24 | 37.04 | 46.58 | E103,Y115 |
|  | 5 | A0A0U3AA44 | 0.21 | 48.0 | 43.23 | E110,Y122 |
| GO:0008106 | 1 | A0A2K6B8V5 | 0.72 | 3.7 | 36.67 | E210,C244 |
|  | 2 | O13848 | 0.19 | 54.55 | 41.2 | Y54,H109 |
|  | 3 | O13848 | 0.36 | 59.09 | 42.25 | Y54,H109 |
|  | 4 | A0A2K5NL52 | 0.55 | 3.57 | 39.67 | E210,C244 |
|  | 5 | P30838 | 0.76 | 10.0 | 35.67 | E210,C244 |
| GO:0008234 | 1 | P05993 | 0.15 | 68.18 | 51.04 | H31,N58 |
|  | 2 | P09648 | 0.17 | 74.19 | 54.59 | C25,H165,N185 |
|  | 3 | P83443 | 0.22 | 64.71 | 42.25 | C26,H159,N176 |
|  | 4 | P83443 | 0.17 | 70.97 | 46.95 | C26,H159,N176 |
|  | 5 | A5YVK8 | 0.2 | 76.47 | 51.63 | C14,H146,N162 |
| GO:0015035 | 1 | A0A520C1H0 | 0.08 | 66.67 | 57.32 | C7,C10 |
|  | 2 | A0A662FLU1 | 0.15 | 87.5 | 66.67 | C30,C33 |
|  | 3 | A0A3D4FEA4 | 0.21 | 73.33 | 58.54 | C7,C10 |
|  | 4 | A0A2K5CLF8 | 0.07 | 57.14 | 61.46 | C23,C26 |
|  | 5 | A0A7Y2VGP6 | 0.1 | 66.67 | 56.47 | C11,C14 |
| GO:0015036 | 1 | A0A182YSM0 | 0.07 | 66.67 | 63.95 | C11,C14 |
|  | 2 | A0A1F9KUL7 | 0.1 | 73.33 | 55.56 | C32,C35 |
|  | 3 | A0A7V3TX07 | 0.11 | 68.75 | 54.63 | C32,C35 |
|  | 4 | A0A550H2S0 | 0.1 | 62.5 | 63.16 | C20,C23 |
|  | 5 | A0A3D4FEA4 | 0.1 | 80.0 | 63.41 | C7,C10 |
| GO:0016298 | 1 | A0A851Y268 | 0.23 | 51.61 | 36.67 | S168,D194,H279 |
| GO:0016407 | 1 | A0A4P7JQ37 | 0.24 | 60.0 | 45.33 | E107,Y119 |
|  | 2 | A0A3B9NVM9 | 0.16 | 41.67 | 48.63 | E103,Y115 |
|  | 3 | A0A2W4VE46 | 0.23 | 61.54 | 48.67 | E105,Y117 |
|  | 4 | A0A0P6VE99 | 0.76 | 50.0 | 45.62 | E111,Y123 |
|  | 5 | A0A654L334 | 0.26 | 46.15 | 41.72 | E99,Y111 |
| GO:0016408 | 1 | Q7A1P9 | 0.15 | 61.9 | 43.33 | C86,H338 |
| GO:0016410 | 1 | D4XNE3 | 0.2 | 76.0 | 44.44 | E99,Y111 |
|  | 2 | A0A6N6RS47 | 0.72 | 36.0 | 44.59 | E104,Y116 |
|  | 3 | A0A2L1WHT7 | 0.25 | 52.0 | 48.67 | E105,Y116 |
|  | 4 | A0A6N1N5Z4 | 0.19 | 64.0 | 42.76 | E99,Y111 |
|  | 5 | A0A7C5L4R9 | 0.65 | 33.33 | 41.96 | E97,Y109 |
| GO:0016419 | 1 | A0A3B0J3W0 | 0.18 | 53.85 | 37.0 | S92,H201 |
|  | 2 | A0A077PZH2 | 0.17 | 46.43 | 41.67 | S92,H201 |
|  | 3 | A0A0K2ART5 | 0.24 | 61.54 | 32.67 | S89,H193 |
|  | 4 | A0A416WG28 | 0.27 | 53.85 | 30.67 | S87,H198 |
|  | 5 | A0A3D3W9N5 | 0.28 | 46.15 | 35.0 | S90,H195 |
| GO:0016420 | 1 | A0A2D4TSP9 | 0.08 | 57.14 | 36.0 | S90,H198 |
|  | 2 | A0A1C6NL35 | 0.11 | 56.0 | 36.67 | S88,H193 |
|  | 3 | A0A536FEV0 | 0.27 | 48.15 | 43.0 | S114,H221 |
|  | 4 | H2IQE4 | 0.3 | 52.0 | 23.33 | S86,H195 |
|  | 5 | A0A101C5L5 | 0.22 | 46.15 | 33.67 | S87,H195 |
| GO:0016620 | 1 | A0A7C7GGH8 | 0.14 | 66.67 | 31.0 | E161,C195 |
|  | 2 | A0A3D0RY98 | 0.12 | 58.82 | 40.67 | E232,C266 |
|  | 3 | A0A511J2Y8 | 0.16 | 66.67 | 46.33 | E253,C287 |
|  | 4 | A0A1M5N6Y7 | 0.22 | 61.54 | 48.33 | E233,C267 |
|  | 5 | A0A1R3FFM7 | 0.21 | 70.37 | 41.33 | E217,C251 |
| GO:0016667 | 1 | A0A522QSK5 | 0.27 | 10.53 | 51.82 | C32,C35 |
|  | 2 | A0A352HRW1 | 0.04 | 62.5 | 59.3 | C7,C10 |
|  | 3 | A0A3D4FEA4 | 0.08 | 47.06 | 54.88 | C7,C10 |
|  | 4 | A0A352PK77 | 0.1 | 80.0 | 51.92 | C30,C33 |
|  | 5 | A0A3C0KDU2 | 0.04 | 50.0 | 70.15 | C34,C37 |
| GO:0016855 | 1 | A0A256BFD5 | 0.28 | 19.05 | 30.43 | C78,C187 |
| GO:0016864 | 1 | Q0E0I1 | 0.09 | 90.0 | 39.67 | C75,C78 |
| GO:0016884 | 1 | E0NNP0 | 0.21 | 64.52 | 44.33 | K69,S144,S168 |
| GO:0030060 | 1 | K0J107 | 0.28 | 55.0 | 40.33 | D173,H200 |
|  | 2 | K0J107 | 0.23 | 76.19 | 40.33 | D173,H200 |
|  | 3 | K0J107 | 0.36 | 52.38 | 40.0 | D173,H200 |
|  | 4 | K0J107 | 0.32 | 63.64 | 39.0 | D173,H200 |
| GO:0032296 | 1 | A0A149VUG9 | 0.21 | 50.0 | 37.83 | D47,E119 |
|  | 2 | Q2NB81 | 0.18 | 50.0 | 39.01 | D48,E119 |
| GO:0033818 | 1 | A0A535CXZ2 | 0.24 | 65.62 | 44.0 | C129,H260,N290 |
|  | 2 | A0A0J9FDV5 | 0.25 | 54.55 | 32.33 | C116,H256,N286 |
|  | 3 | A0A2V5ZBS8 | 0.59 | 33.33 | 35.67 | C127,H267,N297 |
|  | 4 | A0A2V3W4S1 | 0.53 | 37.5 | 32.0 | C112,H237,N267 |
|  | 5 | A0A6I3ZG68 | 0.61 | 35.29 | 40.33 | C122,H258,N289 |
| GO:0047661 | 1 | A0A252DYH1 | 0.23 | 25.0 | 26.62 | C91,C200 |
|  | 2 | A0A0R2B670 | 0.36 | 56.0 | 40.21 | C73,C184 |
|  | 3 | A0A5M8R285 | 0.26 | 52.0 | 25.28 | C75,C188 |
|  | 4 | R7IW21 | 0.58 | 31.82 | 25.9 | C79,C190 |
| GO:0051920 | 1 | A0A3P1CVP9 | 0.12 | 26.67 | 33.67 | C158,C161 |
|  | 2 | A0A243RTW8 | 0.15 | 53.33 | 41.57 | C131,C134 |
| GO:0052689 | 1 | A0A4Y8L3A9 | 0.32 | 32.26 | 28.0 | S93,D192,H222 |
|  | 2 | A0A2E9H3J6 | 0.25 | 41.94 | 32.55 | S81,D206,H234 |
|  | 3 | A0A4R0ZJD2 | 0.22 | 35.71 | 29.48 | S98,D194,H223 |
|  | 4 | A0A7X1TML9 | 0.35 | 44.83 | 30.33 | S186,D277,H306 |
|  | 5 | A0A523I7I3 | 0.32 | 44.83 | 27.0 | S169,D255,H285 |
| GO:0052745 | 1 | Q3ZCK3 | 0.13 | 47.62 | 34.0 | D51,T122 |
|  | 2 | Q9Z0S1 | 0.64 | 33.33 | 32.0 | D51,T122 |
| GO:0070001 | 1 | A0A1Y1ZC50 | 0.09 | 60.87 | 33.33 | D17,D198 |
|  | 2 | A0A818CU57 | 0.12 | 69.57 | 48.0 | D304,D485 |
|  | 3 | A0A3B5Y7I3 | 0.49 | 59.09 | 35.33 | D126,D349 |
|  | 4 | F9XJD5 | 0.2 | 56.52 | 32.0 | D91,D273 |
|  | 5 | A0A667INK2 | 0.12 | 60.87 | 35.67 | D94,D281 |
| GO:0070008 | 1 | Q8L9Y0 | 0.22 | 67.86 | 48.67 | S185,D395,H447 |
|  | 2 | Q9CAU2 | 0.49 | 46.67 | 32.33 | S183,D363,H416 |
|  | 3 | Q869Q8 | 0.34 | 58.62 | 38.33 | S233,D414,H474 |
| GO:0101005 | 1 | A6NNY8 | 0.14 | 71.43 | 42.33 | C87,H380 |
|  | 2 | Q9LEW0 | 0.14 | 71.43 | 45.33 | C186,H491 |
|  | 3 | P62068 | 0.15 | 65.0 | 42.67 | C44,H313 |
|  | 4 | Q52KZ6 | 0.13 | 66.67 | 43.33 | C48,H317 |
|  | 5 | Q9FPS3 | 0.17 | 61.11 | 45.67 | C206,H510 |

